# Two complete genomes of male-killing *Wolbachia* infecting *Ostrinia* moth species illuminate their evolutionary dynamics and association with hosts

**DOI:** 10.1101/2022.05.08.491107

**Authors:** Tomohiro Muro, Hiroyuki Hikida, Takeshi Fujii, Takashi Kiuchi, Susumu Katsuma

## Abstract

*Wolbachia* is an extremely widespread endocellular symbiont which causes reproductive manipulation on various arthropod hosts. Male progenies are killed in *Wolbachia*-infected lineages of the Japanese *Ostrinia* moth population. While the mechanism of male killing and the evolutionary interaction between host and symbiont are significant concerns for this system, the absence of *Wolbachia* genomic information has limited approaches to these issues. We determined the complete genome sequences of *w*Fur and *w*Sca, the male-killing *Wolbachia* of *O. furnacalis* and *O. scapulalis*. The two genomes shared an extremely high degree of homology, with over 95% of the predicted protein sequences being identical. A comparison of these two genomes revealed nearly minimal genome evolution, with a strong emphasis on the frequent genome rearrangements and the rapid evolution of ankyrin repeat-containing proteins. Additionally, we determined the mitochondrial genomes of both species’ infected lineages and performed phylogenetic analyses to deduce the evolutionary dynamics of *Wolbachia* infection in the *Ostrinia* clade. According to the inferred phylogenetic relationship, *Wolbachia* infection was established in the *Ostrinia* clade prior to the speciation of related species such as *O. furnacalis* and *O. scapulalis*. Simultaneously, the relatively high homology of mitochondrial genomes suggested recent *Wolbachia* introgression between infected *Ostrinia* species. The findings of this study collectively shed light on the host-symbiont interaction from an evolutionary standpoint.

**Significance:** Despite the growing number of publicly available *Wolbachia* genome sequences, only a few high-quality male-killer genomes exist, particularly those found in lepidopteran hosts. The complete genomes of two male-killing *Wolbachia* of *Ostrinia* moth hosts were determined in this study. The genomic data obtained here will be used to elucidate the mechanism of reproductive manipulation and the origins of this endosymbiont’s extraordinary diversity. Additionally, phylogenetic analysis of mitochondria and *Wolbachia* revealed the evolutionary history of *Ostrinia* hosts and *Wolbachia*. The inferred dynamic pattern of infection adds to our understanding of evolution and ecology of *Wolbachia* endosymbiont, a promising agent for biological pest control.

## Introduction

Endosymbionts have a variety of effects on host biology. While mutual symbionts are necessary for the growth of their hosts, there are numerous instances in which symbionts drastically impose parasitic phenotypes on their hosts (Goryacheva & Andrianov 2021). *Wolbachia* is one of these bacteria, notable for its widespread distribution among insects (Weinert et al. 2015). *Wolbachia* infection has been reported in a wide variety of insect lineages, as well as some other terrestrial arthropods and filarial nematodes. *Wolbachia* strains are classified into at least 17 supergroups, in which Supergroups A and B contain a majority of strains found in arthropod taxa (Kaur et al. 2021). In many associations between *Wolbachia* and its hosts, *Wolbachia* induces reproductive manipulation on the hosts, which typically results in infected females having a higher fitness level than uninfected females (Goryacheva & Andrianov 2021; Kaur et al. 2021). Due to the vertical transmission of *Wolbachia* from mothers to their offspring, such manipulation is thought to spread *Wolbachia* infection throughout the host population. *Wolbachia* has four modes of reproductive manipulation: cytoplasmic incompatibility, feminization of genetic males, induction of parthenogenesis, and male killing (Kaur et al. 2021). Male killing is the process by which male offspring of infected mothers are killed during embryonic development or at a later stage. Although male killing has been observed in a variety of bacterial parasites, including at least four different groups of bacteria (Goryacheva & Andrianov 2021), *Wolbachia*-induced male killing has ever been reported for relatively limited groups, including *Drosophila* flies (Dyer & Jaenike 2004; Hurst et al. 2000; Sheeley & McAllister 2009), ladybugs (Elnagdy et al. 2013; Hurst et al. 1999) and some lepidopteran insects such as *Acraea* species (Hurst et al. 1999; Jiggins et al. 1998, 2000), *Hypolimnas bolina* (Dyson et al. 2002), *Homona magnanima* (Arai et al. 2020), and *Ostrinia* species (Fukui et al. 2015; Kageyama & Traut 2004).

Several agricultural pests are found in the genus *Ostrinia*, which is a member of the Crambidae family of Lepidoptera. Historically, the genus was largely classified into three groups and are denoted by the letters I, II, and III, according to the morphology of male genitalia (Mutuura & Munroe 1970). However, a recent phylogenomic study confirmed the monophyly of only group III and proposed a new classification system in which the genus is divided into three new species groups (Clade I, II, and III) (Yang et al. 2021). All species of group III (Mutuura & Munroe, 1970) were incorporated into Clade III, the *Ostrinia nubilalis* species group, along with some previously classified species in group II. Both *O. furnacalis* (Asian corn borer) and *O. scapulalis* (Adzuki bean borer), classified as group III or the new Clade III, are important agricultural pests. *O. furnacalis* is a significant maize pest in Asia, causing substantial economic damage, whereas *O. scapulalis* attacks some crops such as hops, hemp and legumes (Ishikawa et al. 1999; Yang et al. 2021).

*Wolbachia* causes male killing in Japanese populations of both *O. furnacalis* and *O. scapulalis* (Fukui et al. 2015; Kageyama & Traut 2004). The infection rate of *Wolbachia* in *O. furnacalis* is comparatively low: a previous survey reported that 13 individuals out of 79 collected females were infected (Kageyama et al. 2002). The striking feature of infected strains of *Ostrinia* moths is that when the infection is cured with antibiotics, cured females produce all-male broods due to female-specific death (Kageyama & Traut 2004). This is in contrast to the situation with other male-killing *Wolbachia*, where antibiotic treatment restores a balanced sex ratio encompassing both sexes, as observed in *Acraea* butterflies (Jiggins et al. 1998, 2000), *Hypolimnas bolina* (Charlat et al. 2007), and *Homona magnanima* (Arai et al. 2020). When *Wolbachia* is present in *O*. *furnacalis*, the expression of *Ostrinia furnacalis Masculinizer* gene (*OfMasc*) is downregulated at the embryonic stage (Fukui et al. 2015). *OfMasc* is an ortholog of *Masculinizer* gene, which was first discovered in the silkworm *Bombyx mori* and demonstrated to be required for both masculinization and dosage compensation (Kiuchi et al. 2014). *OfMasc* is thought to have masculinizing activity in *O. furnacalis* because knockdown of *OfMasc* results in the expression of female-type splicing variants of *Ostrinia furnacalis doublesex* (*Ofdsx*) in male embryos (Fukui et al. 2018). Males infected with *Wolbachia* were rescued from male killing by the injection of *OfMasc* cRNA, and the expression of Z-linked genes was increased in *Wolbachia*-infected embryos (Fukui et al. 2015). Therefore, the depletion of *OfMasc*, which probably results in the failure of dosage compensation, may be the underlying mechanism of male-specific death in *Wolbachia* infection. Female-specific death in antibiotic treatment is also attributed to the aberrant expression of sex chromosome-linked genes (Sugimoto et al. 2015). These studies highlight the *Wolbachia*’s ingenious strategy for reproductive manipulation of *Ostrinia* hosts, generating increased interests in the evolutionary context of their association.

Genomic information is a critical resource for characterizing host-symbiont interactions. Recent advances in sequencing technology have resulted in the accumulation of *Wolbachia* genome sequences, which have aided in the identification of factors responsible for reproductive manipulation (Le Page et al. 2017; Perlmutter et al. 2019) as well as the characterization of host-symbiont association dynamics at an ecological scale (Cooper et al. 2019; Turelli et al. 2018; Wolfe et al. 2021). In order to obtain reliable insights, high-quality genome sequences from a diverse set of *Wolbachia* strains representing both phenotypic and phylogenetic diversity are required. However, there are a few genome available for male-killing *Wolbachia* (Duplouy et al. 2013; Hill et al. 2021; Metcalf et al. 2014). One is the complete genome sequence of *w*Inn strain (Supergroup A), a male-killer of *Drosophila innubila* (Hill et al. 2021). The other two are draft genome assemblies. *w*Rec strain (Supergroup A), which causes cytoplasmic incompatibility in its native host *D. recens* and male killing in an introgressed host *D. subquinaria*, has the genome with reduced prophage regions compared to its close relative *w*Mel strain (Metcalf et al. 2014). The only Lepidoptera-associated male-killing strain with a draft genome is *w*Bol1-b, a Supergroup B strain that infects *H. bolina* butterfly. The *w*Bol1-b genome assembly comprised 144 contigs, 91 of which were organized into 13 scaffolds (Duplouy et al. 2013). Along with the scarcity of genomic data on male-killing *Wolbachia*, the number of *Wolbachia* genome assemblies infecting Lepidopteran hosts is also limited. Indeed, no complete genomes of *Wolbachia* associated with Lepidoptera have been published, although a few are available in public databases. Despite the low number of Lepidoptera-associated *Wolbachia* genomes, a significant number of *Wolbachia* strains are present in Lepidoptera, presumably at a higher rate than in other arthropods (Ahmed, Araujo-Jnr, et al. 2015; Ilinsky & Kosterin 2017).

Mitochondrial genomes are also useful for deciphering host-symbiont associations. Since both inherited symbionts and mitochondria are transmitted maternally in infected lineages, examining mitochondrial haplotypes and *Wolbachia* infection status enables us to infer the mode of *Wolbachia* acquisition among allied species (Baldo et al. 2008; Charlat et al. 2009; Cooper et al. 2019; Sucháčková Bartoňová et al. 2021). Specifically, there are three distinct ways in which *Wolbachia* can be acquired by host species: cladogenic acquisition, introgression, and horizontal transmission (Cooper et al. 2019; Raychoudhury et al. 2009; Turelli et al. 2018). Cladogenic acquisition occurs when related species codiverged (along cladogenesis) with *Wolbachia* derived from a common ancestor that was previously infected. Alternatively, *Wolbachia* infection can be introduced into a species via hybridization with infected sibling species and subsequent backcrossing. This is referred to as introgression. The relatively low level of genetic divergence of mitochondria and *Wolbachia* between host species can serve as an indicator in this case (Meany et al. 2019; Raychoudhury et al. 2009). The third mode is horizontal transmission, which typically results in discordance of phylogenetic relationship between *Wolbachia* and the host’s mitochondria.

Here, we report the complete genome sequences of male-killing *Wolbachia* infecting *O. furnacalis* and *O. scapulalis* (referred to as *w*Fur and *w*Sca, respectively), as well as the mitochondrial genomes of both hosts of infected lines. The comparison of these male-killing *Wolbachia* genomes revealed a trend in *Wolbachia* genome evolution. Along with mitochondrial genome analysis, the obtained genomic data enabled us to infer the evolutionary history of the *Wolbachia* infection in *Ostrinia* moths.

## Materials & Methods

### Insects

*w*Fur-infected *O. furnacalis* and *w*Sca-infected *O. scapulalis* were used in this study. An *O. furnacalis* line infected with *w*Fur was previously described (Fukui et al. 2015) and is maintained in our laboratory. *w*Sca-infected *O. scapulalis* line was newly established by collecting an infected female moth in Matsudo, Japan (35.8° N, 139.9° E) in September 2020. Sex pheromone analysis was used to determine the species of collected *Ostrinia* females. Pheromone glands (PG) of three females from each matriline were immersed in hexane to extract sex pheromone. The PG extracts were analyzed using a GC-MS unit (QP2010 SE GC-MS, Shimadzu) equipped with a capillary column (DB-Wax, 0.25 mm i.d. × 30 m; Agilent Technologies, Santa Clara, CA, USA). The initial column oven temperature of 80°C was held for 2 min and then increased at 8°C/min to 240°C. The final temperature was maintained for 2 min. The flow rate of the helium carrier gas was 1.0 mL/min. The diagnostic ion of the acetate ion at 61 (*m*/*z*) and its eliminated formation at 194 (*m*/*z*) were used for GC-MS analysis, in addition to the retention time. Both *O. furnacalis* and *O. scapulalis* lines were reared on an artificial diet (Insecta-LFS, Nosan Corp.) at 25°C under a photoperiod of 16L8D and maintained by crossing with males of *Wolbachia*-uninfected counterpart lines derived from females collected in Nishi-Tokyo, Japan (35.7° N, 139.5° E). The Infection was treated by adding 0.3 g tetracycline-HCl to a 500 g artificial diet.

### Reverse transcription-polymerase chain reaction (RT-PCR)

RT-PCR was used to examine the sex-specific splicing patterns of *O. scapulalis doublesex* (*Osdsx*). Egg masses (96 hours post oviposition; hpo) from infected, uninfected, and cured lines were sampled. Total RNA and genomic DNA were prepared simultaneously from the egg masses using TRI Reagent (Molecular Research Center, Inc.) according to the manufacturer’s protocol. Total RNA was subjected to reverse transcription using AMV-RTase and an oligo-dT primer (TaKaRa). The resultant cDNA was used to perform PCR to examine the sex-specific splicing patterns of *Osdsx*. Genomic PCR of *Wolbachia*-specific *wsp* gene was used to examine the infection status. PCR was performed with KOD FX Neo (TOYOBO) for *Osdsx* and KOD One (TOYOBO) for *wsp*. Primer sequences were reported previously (Sugimoto et al. 2010).

### Molecular sexing

*Ostrinia* larvae were sexed using a previously reported method for *O. furnacalis* (Fukui et al. 2015). Briefly, genomic DNA was purified using DNeasy Blood & Tissue Kit (QIAGEN), and the dosage of Z-linked genes (*Tpi* and *kettin*) normalized by the autosome-linked gene (*EF-1α*) was evaluated by genomic qPCR. Individual larvae were photographed prior to DNA extraction, and body lengths were measured using ImageJ software.

### Genome sequencing and assembly

Genomic DNA was purified from the ovaries of *w*Fur-infected *O. furnacalis* and *w*Sca-infected *O. scapulalis* females using Genomic-tip 100/G (QIAGEN) according to the manufacturer’s protocol. Prepared DNA samples were subjected to genome sequencing on NovaSeq 6000 (Illumina) and Sequel (PacBio) platforms. For *O. furnacalis*, two DNA samples were prepared each from an infected female. One of them was used for Illumina sequencing, and subsequently the remainder was combined with the other sample for PacBio (Sequel system) sequencing. For *O. scapulalis*, one and four ovaries were used to prepare DNA samples for Illumina and PacBio (Sequel II system) sequencing, respectively. Long-reads obtained by PacBio sequencing were assembled using Canu v2.1 assembler (Koren et al. 2017). An estimated genome size of 438.5 Mb was given as a parameter with reference to the deposited genome assembly of *O. furnacalis* (accession number: GCA_004193835.1).

### Construction of *Wolbachia* genomes

Both *w*Fur and *w*Sca genomes were constructed through a following procedure. To identify contigs derived from *Wolbachia* in Canu assembly, the generated contigs were subjected to blastn search as a query against RefSeq representative prokaryotic genomes database (ref_prok_rep_genomes) with the following parameters: “evalue 1e-20”, “max_target_seqs 60”, and “max_hsps 10”. Illumina short-reads were mapped to the draft *Wolbachia* contig with BWA-MEM v.0.7.17 (Li 2013), and the resultant SAM file was converted to BAM format using samtools v.1.9 (Danecek et al. 2021). Pilon v.1.23 (Walker et al. 2014) was utilized to polish the draft *Wolbachia* contig. In order to locate the overlap region at the beginning and end of the sequence, a blastn search was performed using the same polished contig as both a subject and a query. The overlap region was manually modified to construct a single-coverage genome (see Results). The oriC regions of *w*Fur and *w*Sca were identified by performing a homology search against the oriC sequence of *w*Mel strain. The oriC sequence of *w*Mel strain (DoriC ID: ORI10030016) was retrieved from the DoriC 5.0 database (Gao et al. 2013). The binding sites in the oriC sequence were searched using the same criteria as previously described (Ioannidis et al. 2007). For genome annotation, stand-alone PGAP v5.2 (2021-05-19.build5429) (Tatusova et al. 2016) was utilized.

### Construction of mitochondrial genomes

By performing a blastn search on mitochondrial gene sequences in the Canu assembly of *w*Fur- infected *O. furnacalis*, a contig corresponding to a mitochondrial genome (mitogenome) was identified. Publicly available *COI* (Gene ID 65331444), *COII* (65331445), and *ND5* (65331454) sequences from an *O. furnacalis* mitogenome (accession number: NC_056248.1) were used as queries. Illumina short-reads were mapped to the identified mitogenome contig with BWA-MEM v.0.7.17 (Li 2013). Visual inspection of mapped reads was performed using IGV v.2.8.7 (Thorvaldsdóttir et al. 2013), and overlaps were manually modified to construct a single-coverage mitogenome. Subsequently, Illumina short-reads from *w*Sca-infected *O. scapulalis* were mapped to the constructed mitogenome of *w*Fur-infected *O. furnacalis* with BWA-MEM v.0.7.17 and visualized using IGV v.2.8.7. Detected mutations were manually modified to construct a mitogenome of *w*Sca-infected *O. scapulalis*.

### Genome analysis

Some *Wolbachia* genes were searched in the *w*Fur and *w*Sca genomes using tblastn. The deduced amino acid sequences of *w*Mel strain *cifA* (AAS14330.1), *cifB* (AAS14331.1), *wmk* (AAS14326.1), and *TomO* (AAS14922.1) were downloaded from GenBank and used as queries. To calculate the average nucleotide identity (ANI) between *Wolbachia* or mitochondrial genomes, FastANI v1.33 (Jain et al. 2018) was used. In order to conduct syntenic analysis, the *w*Fur and *w*Sca genomes were aligned using MUMmer v4.0.0rc1 (Marçais et al. 2018) in the “nucmer” mode, and a dot plot was generated with the packaged program, mummerplot. A comparison of the gene repertoires of *w*Fur and *w*Sca was performed. To obtain non-redundant sequences for each strain’s annotated protein sequences, duplicated identical sequences were removed using SeqKit (Shen et al. 2016) in “rmdup” mode. Reciprocal blastp searches were used to identify orthologous gene pairs between the non-redundant *w*Fur and *w*Sca protein sequences. The hmmscan program was used in conjunction with HmmerWeb v.2.41.2 (Potter et al. 2018) to conduct a domain search against the Pfam 35.0 database using an E-value threshold of 0.05.

### Phylogenetic analysis

For *Wolbachia*, annotated protein sequences derived from 136 *Wolbachia* genome assemblies found in NCBI RefSeq as of December 2021 were downloaded. Additionally, two complete genome assemblies included in the analysis were not registered in RefSeq but were found in GenBank (GCA_000953315 and GCA_020995475). BUSCO v.5.2.2 (Simão et al. 2015) was used to determine the completeness of each genome assembly using rickettsiales_odb10. Two assemblies (GCF_000174095.1 and GCF_000167475.1) with low complete BUSCO percentage were excluded from the subsequent analysis. Using OrthoFinder v2.5.4 (Emms & Kelly 2018) with the following options: “-M msa” and “-os”, single-copy orthologs were identified across the aforementioned assemblies and the *w*Fur and *w*Sca genomes (140 in total; Supplementary table S1). Each of the 63 predicted single-copy orthologous groups was aligned using MAFFT v.7.490 (Katoh & Standley 2013) with default parameters, and the resultant alignments were trimmed using trimAl v.1.4.1 (Capella-Gutiérrez et al. 2009) in the “automated1” mode. The trimmed alignments for each strain were concatenated using SeqKit v2.1.0 (Shen et al. 2016), and were used for phylogenetic analysis.

**Table 1.**
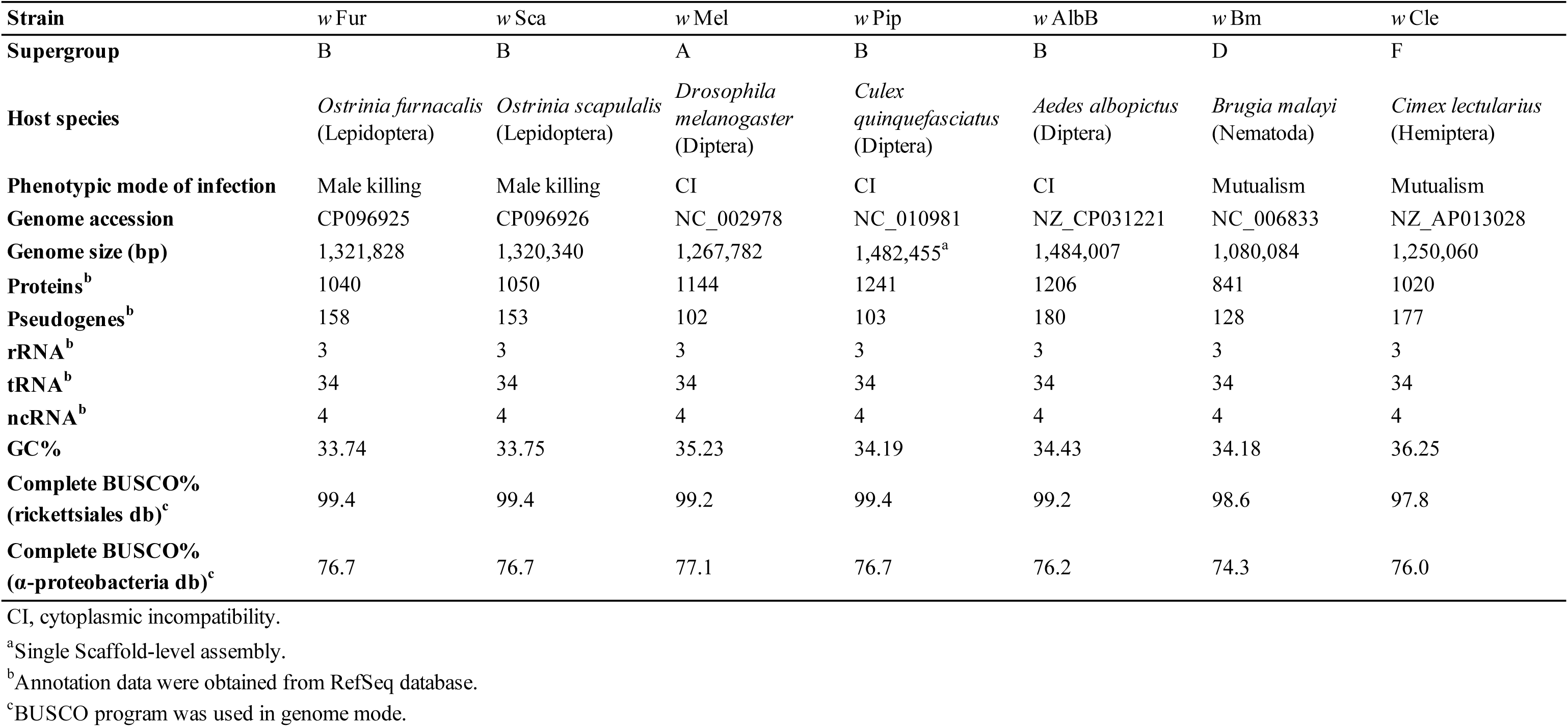
General characteristics of *w* Fur, *w* Sca, and representative *Wolbachia* genomes.

| Strain | *w* Fur | *w* Sca | *w* Mel | *w* Pip | *w* AlbB | *w* Bm | *w* Cle |
| --- | --- | --- | --- | --- | --- | --- | --- |
| **Supergroup** | B | B | A | B | B | D | F |
| **Host species** | *Ostrinia furnacalis*<br>(Lepidoptera) | *Ostrinia scapularis*<br>(Lepidoptera) | *Drosophila melanogaster*<br>(Diptera) | *Culex quinquefasciatus*<br>(Diptera) | *Aedes albopictus*<br>(Diptera) | *Brugia malayi*<br>(Nematoda) | *Cimex lectularius*<br>(Hemiptera) |
| **Phenotypic mode of infection** | Male killing | Male killing | CI | CI | CI | Mutualism | Mutualism |
| **Genome accession** | CP096925 | CP096926 | NC_002978 | NC_010981 | NZ_CP031221 | NC_006833 | NZ_AP013028 |
| **Genome size (bp)** | 1,321,828 | 1,320,340 | 1,267,782 | 1,482,455<sup>a</sup> | 1,484,007 | 1,080,084 | 1,250,060 |
| **Proteins<sup>b</sup>** | 1040 | 1050 | 1144 | 1241 | 1206 | 841 | 1020 |
| **Pseudogenes<sup>b</sup>** | 158 | 153 | 102 | 103 | 180 | 128 | 177 |
| **rRNA<sup>b</sup>** | 3 | 3 | 3 | 3 | 3 | 3 | 3 |
| **tRNA<sup>b</sup>** | 34 | 34 | 34 | 34 | 34 | 34 | 34 |
| **ncRNA<sup>b</sup>** | 4 | 4 | 4 | 4 | 4 | 4 | 4 |
| **GC%** | 33.74 | 33.75 | 35.23 | 34.19 | 34.43 | 34.18 | 36.25 |
| **Complete BUSCO%**<br>**(rickettsiales db)<sup>c</sup>** | 99.4 | 99.4 | 99.2 | 99.4 | 99.2 | 98.6 | 97.8 |
| **Complete BUSCO%**<br>**(α-proteobacteria db)<sup>c</sup>** | 76.7 | 76.7 | 77.1 | 76.7 | 76.2 | 74.3 | 76.0 |

CI, cytoplasmic incompatibility.
<sup>a</sup>Single Scaffold-level assembly.
<sup>b</sup>Annotation data were obtained from RefSeq database.
<sup>c</sup>BUSCO program was used in genome mode.

A maximum likelihood (ML) tree was constructed using IQ-TREE v.1.6.12 (Nguyen et al. 2015), and node support was estimated with 1000 ultrafast bootstrap replicates (Hoang et al. 2018). ModelFinder (Kalyaanamoorthy et al. 2017) was used to determine the best-fitting substitution model, and the HIVw+F+R4 model was chosen. Additionally, we conducted phylogenetic analysis using only *Wolbachia* strains belonging to Supergroup B. *Wolbachia* genomes belonging to Supergroup B were chosen based on the constructed ML tree of all genome assemblies, and subjected to the following analysis. *w*Mel (GCF_000008025.1) and *w*Ri (GCF_000022285.1) genomes (Supergroup A) were also included as outgroups. The phylogenetic analysis was performed in the manner described previously. A total of 339 orthologs were used, with the HIVw+F+R3 model being the best-fitting.

In order to conduct phylogenetic analysis on mitochondrial genomes, we used publicly available sequences from GenBank (accession numbers are shown in Fig. 6). Mitogenomes were annotated de novo using MitoZ v2.4 (Meng et al. 2019) in the “annotate” mode with the following parameters: “--genetic_code 5” and “--clade Arthropoda”. Phylogenetic trees were constructed using the amino acid sequences of 13 protein-coding genes (PCGs) and the nucleotide sequences of 13 PCGs, rRNAs, and tRNAs. In both cases, ML trees were constructed using a partitioned model with partitioning scheme selection (“-m MFP+MERGE” option) in IQ-TREE v.1.6.12 (Chernomor et al. 2016). Alignments of each deduced amino acid sequence were generated using MAFFT v.7.471, trimmed with trimAl v1.4.rev22, and subsequently concatenated. Each protein was assigned a partition. For nucleotide sequences, mitogenomes were aligned directly using MAFFT v.7.471, and partitions for each gene were defined based on annotated positions in a single strain (MN793323.1). The optimal partition scheme and substitution model for each partition were selected based on Bayesian information criterion.

## Results

### 1. Characterization of *w*Sca-induced male killing in *O. scapulalis*

We used *w*Fur-infected *O. furnacalis* (Fukui et al. 2015) and *w*Sca-infected *O. scapulalis* lines for genome sequencing. A *w*Sca-infected *O. scapulalis* line was newly established from a field-collected female moth in this study. We used GC-MS to identify species using hexane extract of pheromone glands. The extract of the collected line contained (E)-11- and (Z)-11- tetradecenyl acetates (retention times were 16.87 min and 16.99 min, respectively), confirming that it is *O. scapulalis* (Ishikawa et al. 1999). This *w*Sca-infected line exhibited a nearly entirely female-biased sex ratio through at least 11 generations, indicating the presence of male killing (Fig. 1A and B). Antibiotics treatment of infection resulted in the occurrence of only male progeny. In order to ascertain the period during which males are killed in *w*Sca-infected *O. scapulalis*, molecular sexing using qPCR was performed on 4- and 14-days post-hatch (dph) larvae. As a result, both females and males were detected in 4 dph larvae, but only females were observed in 14 dph larvae (Fig. 1C). In addition, *w*Sca-infected males were significantly smaller than females in 4 dph larvae, unlike uninfected larvae, which showed no difference in body length between sexes (Fig. 1D). The splicing pattern of *Osdsx*, which represents the phenotype of sexual differentiation, was female in the infected line, male in the cured progeny, and both female and male in the uninfected line (Fig. 1E). These observations demonstrate that *w*Sca disrupts the host’s sex determination cascade from embryogenesis, and that *w*Sca-infected males die during the larval stage of *O. scapulalis*.

**Fig. 1.**
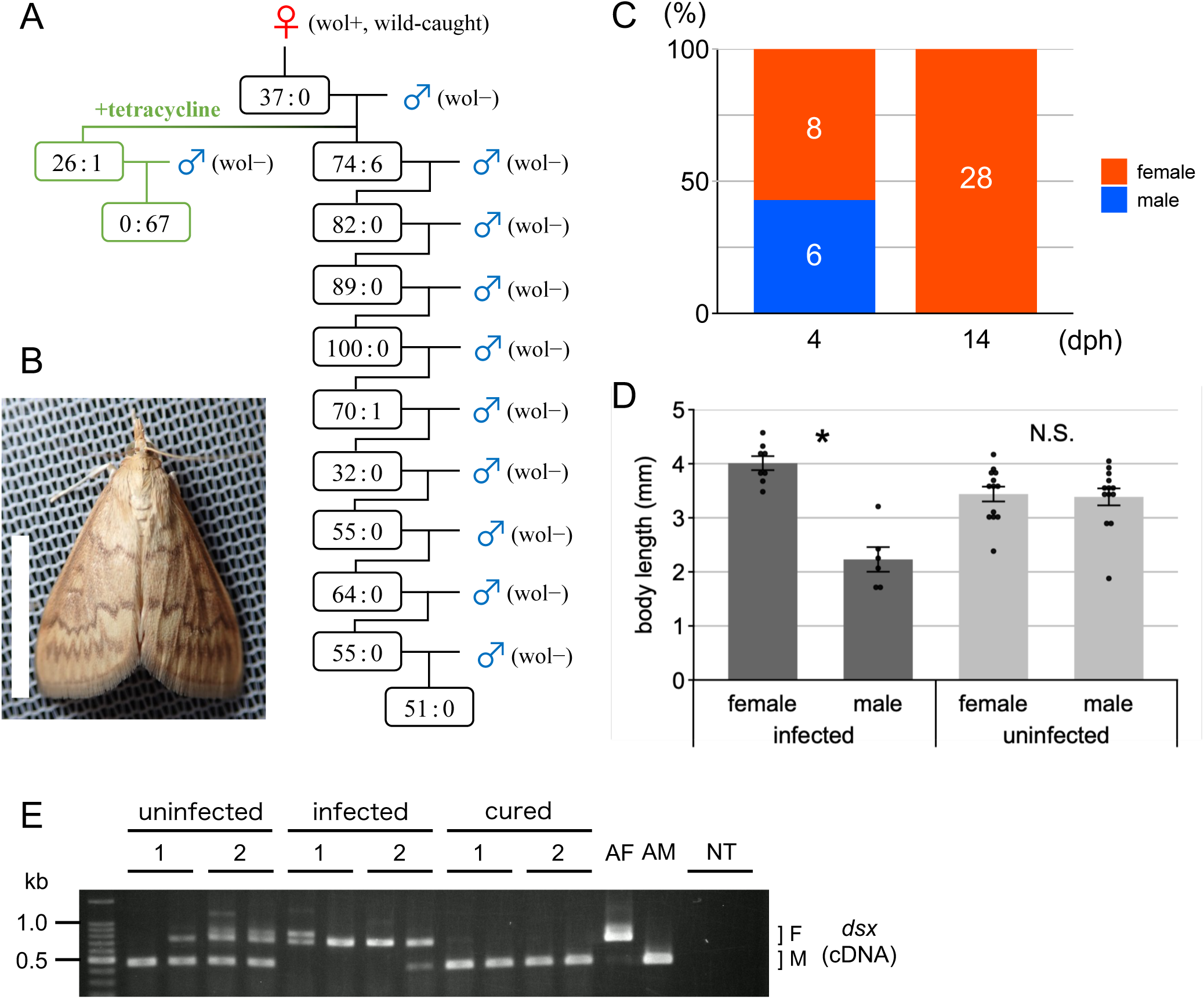
Characterization of male killing in *w*Sca-infected *O. scapulalis*. (A) Brood sex ratios in the *w*Sca-infected matriline. The female:male ratio of each mating is shown. Tetracycline treatment was conducted to remove *w*Sca from a subpopulation of the second generation. Sexing was conducted based on the morphology of the pupal abdominal tips. (B) An adult female moth infected with *w*Sca. Bar, 10 mm. (C) The sex ratios of *w*Sca-infected larvae at 4 and 14 dph. The number indicates the sample size of each group. (D) The body length of *w*Sca- infected and uninfected larvae at 4 dph. Data presented are mean ± standard error of the mean. Dot plots show all data points individually. An asterisk denotes statistical significance (*P* < 0.001; N.S., not significant, *P* > 0.1; Wilcoxon rank-sum test). (E) Splicing patterns of *Osdsx* in uninfected, *w*Sca-infected, and infection-cured embryos at 96 hpo. The results of technical duplicates of the RT-PCR assay are presented for each sample. The numbers indicate individual samples (biological duplicates for each condition). Adult female and male moths were used as positive controls. The letters F and M indicate female- and male-type splicing variants, respectively. AF: adult female; AM: adult male; NT: no template.

### 2. The construction of the *w*Fur and *w*Sca genomes

We conducted both long-read (PacBio) and short-read (Illumina) sequencing. First, long-reads were assembled by Canu assembler (Koren et al. 2017). The resultant assembly of *O. furnacalis* and *O. scapulalis* contained 4,890 and 3,535 contigs, respectively. For each assembly of *O. furnacalis* and *O. scapulalis*, the blastn search using each contig as a query resulted in only one contig that frequently aligned to publicly available *Wolbachia* sequences. Given that both of these two contigs were 1.3–1.4 Mbp in length, which is within the range of typical *Wolbachia* genome sizes (Kaur et al. 2021), and their coverage was among the highest in each assembly (Supplementary Fig. S1), these two contigs were most likely the genomes of *w*Fur and *w*Sca. The polishing process using Illumina reads by Pilon (Walker et al. 2014) identified a few errors in initial contigs (2 changes for *w*Fur and no changes for *w*Sca; all of them were numbers of the same sequential nucleotides, e.g., AAAAA to AAAAAA). Since the polished genomes had an overlap region of approximately 50 kb (*w*Fur) and 80 kb (*w*Sca) at the head and tail of the contigs, duplicated sequences were removed to make the genomes single coverage. When there was inconsistency between the head and tail overlapping regions, we consulted the mapped short-reads to determine which one should be adopted. Due to the circular nature of the *Wolbachia* genome, we manually set the starting points of the sequences to the estimated oriC regions. The oriC region was identified using a blastn search with the *w*Mel oriC sequence as a query. *w*Fur and *w*Sca had identical estimated oriC sequences (405 bp). It contained three DnaA-, four CtrA-, and two IHF-binding sites, and was flanked by *hemE* and CBS domain protein genes, all of which are characteristic of *Wolbachia* (Ioannidis et al. 2007). BUSCO program (Simão et al. 2015) was used to assess the completeness of the genomes. This program was routinely used to determine the presence/absence of benchmarking single-copy orthologs expected to exist among specific taxa. The BUSCO profiles of the *w*Fur and *w*Sca genomes against the Rickettsiales database were identical, with 362/364 single-copy orthologs detected, comparable to the profiles of complete *Wolbachia* genome assemblies (Table 1 and Supplementary Fig. S2). Thus, we ascertained that our assemblies are of sufficient quality to conduct further analyses.

### 3. The properties of the *w*Fur and *w*Sca genomes

The genome of *w*Fur and *w*Sca are 1,321,828 and 1,320,340 bp in length, respectively (Fig. 2). The *w*Fur genome contained 1,040 protein-coding genes, 158 pseudogenes, 34 tRNAs, 3 rRNAs, and 4 ncRNAs, while the *w*Sca genome contained 1,050 protein-coding genes, 153 pseudogenes, 34 tRNAs, 3 rRNAs, and 4 ncRNAs. Some of host-manipulating *Wolbachia* genes were screened individually (Supplementary table S2). Each of the two genomes encodes a putatively pseudogenized copy of *cifA*, one of the two factors responsible for cytoplasmic incompatibility (Le Page et al. 2017). Regarding *cifB*, one gene shares partial homology with the *w*Mel *cifB* sequence, but there do not appear to be any full-length copies. Two intact copies of *wmk*, a candidate gene for male killing (Perlmutter et al. 2019), were found in each of the two genomes. Three genes (including two putative pseudogenes) share homology with *TomO* gene, a growth-inhibitory and RNA-interacting factor of *w*Mel (Ote et al. 2016). However, two of them might be a single ORF misinferred during the automated annotation process.

**Fig. 2.**
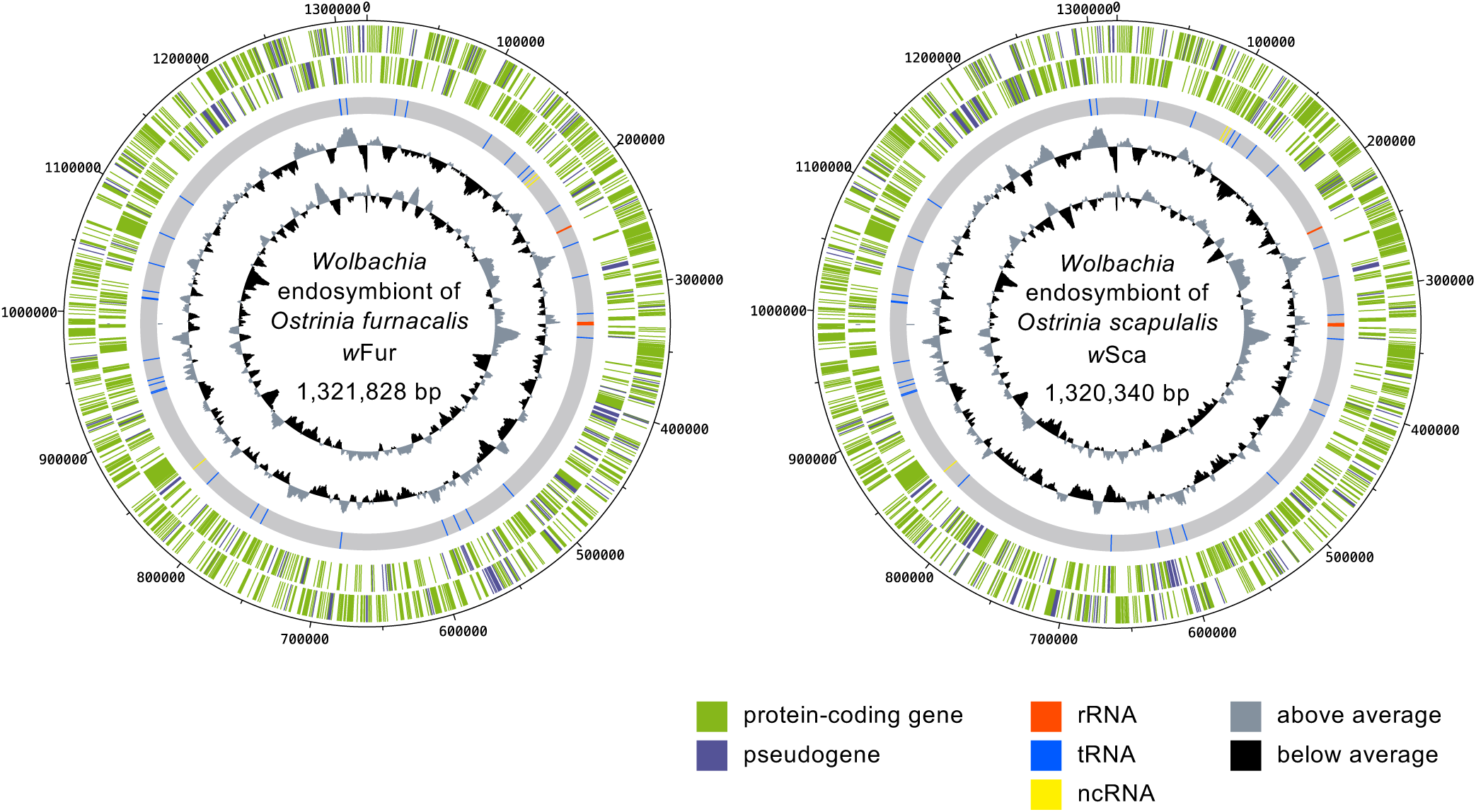
Circular map of the *w*Fur and *w*Sca genomes. Circles arranged in order from outer to inner indicate the following: The locations of annotated coding sequences (CDS) on the positive (outermost) and the negative (second outermost) strands, the position of annotated RNAs, GC content, and GC skew. The GC content and GC skew are calculated in 10 kb windows and expressed as deviations from an average of the whole sequence.

Subsequently, phylogenetic analysis using 138 *Wolbachia* genome assemblies was performed to ascertain the phylogenetic relationship of *w*Fur and *w*Sca within *Wolbachia* (Fig. 3 and Supplementary Fig. S3). As a result, *w*Fur and *w*Sca were assigned to Supergroup B collectively. The most closely related strain was that which infects the Noctuid moth, *Spodoptera picta*. Notably, the second closest strain was *w*Tpre, which is found in a parasitoid wasp *Trichogramma pretiosum* and is responsible for parthenogenesis (Lindsey et al. 2016).

**Fig. 3.**
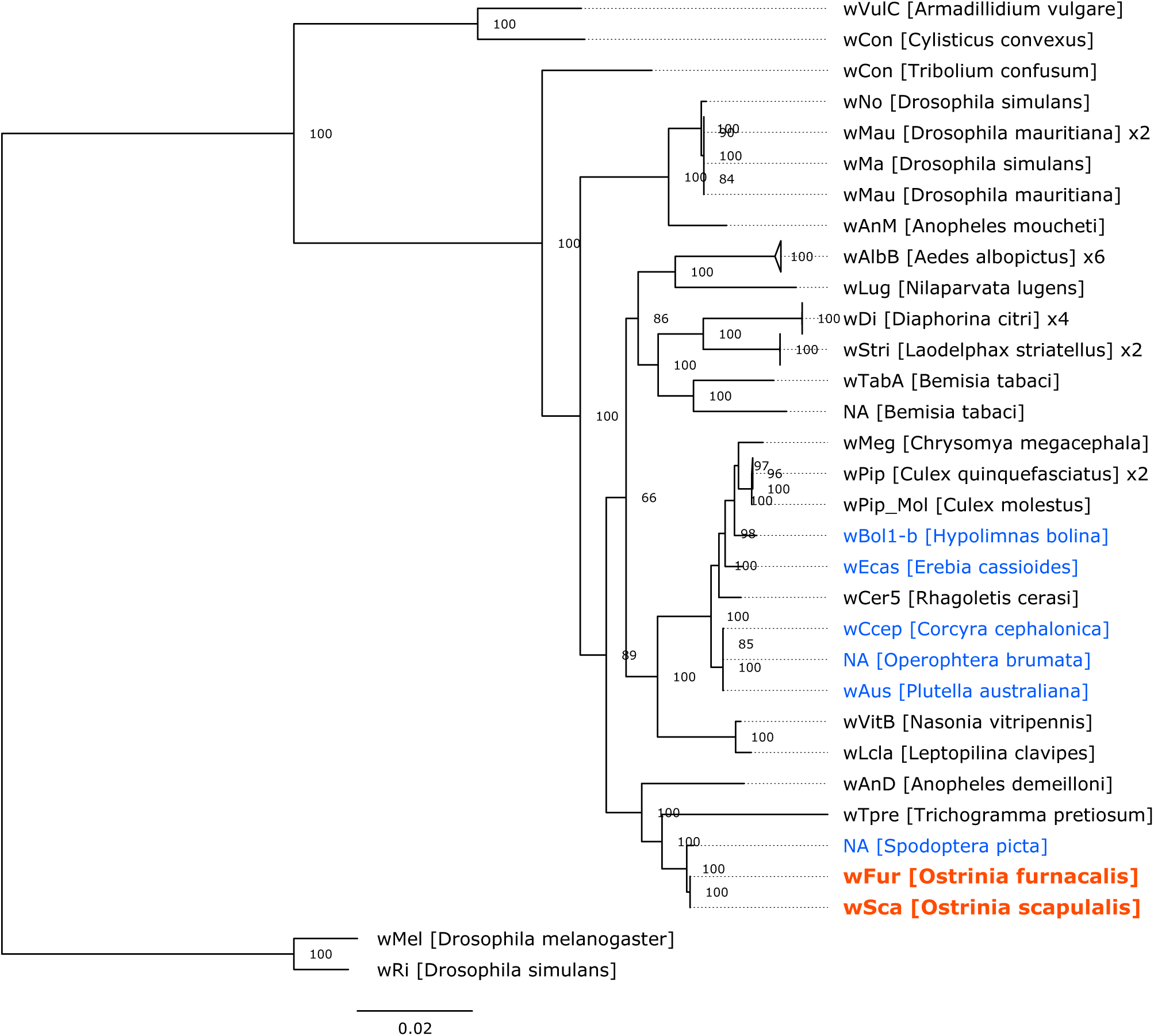
Phylogenetic relationship of Supergroup B *Wolbachia* genomes. The maximum likelihood tree constructed from concatenated protein sequences of 339 single-copy orthologs is shown. *w*Mel and *w*Ri (Supergroup A) were used as outgroups. The strain names and their hosts are labeled. If no suitable strain name is available, it is denoted by “NA”. Clades composed of almost the same strains are collapsed, and the number of contained strains is labeled. Branch support calculated using 1000 replicates of ultrafast bootstrap is shown on the nodes. The sequences determined in this study are highlighted in red and bold, while strains associated with other lepidopterans are in blue.

Intriguingly, the *w*Fur and *w*Sca strains did not cluster phylogenetically with the *w*Bol1-b strain, which induces male killing in the *H. bolina* butterfly (Duplouy et al. 2013).

### 4. Comparative analysis of the *w*Fur and *w*Sca genomes

The ANI calculated by fastANI program (Jain et al. 2018) was 99.9755%, indicating extremely high similarity between these two *Wolbachia*. At the genome structure level, however, we found several large inversions between them (Fig. 4). To compare gene repertoires, we first removed duplicated, identical protein sequences from each genome. We then used blastp to compare the strains’ unique sequences (referred to as “non-redundant sequences”). As a result, 976 out of 982 non-redundant *w*Fur sequences have reciprocal best hit counterparts in 993 non-redundant *w*Sca sequences. Out of these 976 sequences, 952 were identical between two strains. The remaining 24 sequences differed between the strains to a varying degree; whereas 18 have a single amino acid substitution, one of the remaining six had 24 amino acid substitutions and seven gaps (Supplementary table S3). Most of the 6 *w*Fur- and 17 *w*Sca-specific sequences were hypothetical proteins or transposase variants (Supplementary table S3). However, *w*Sca uniquely has three ankyrin repeat-containing proteins.

**Fig. 4.**
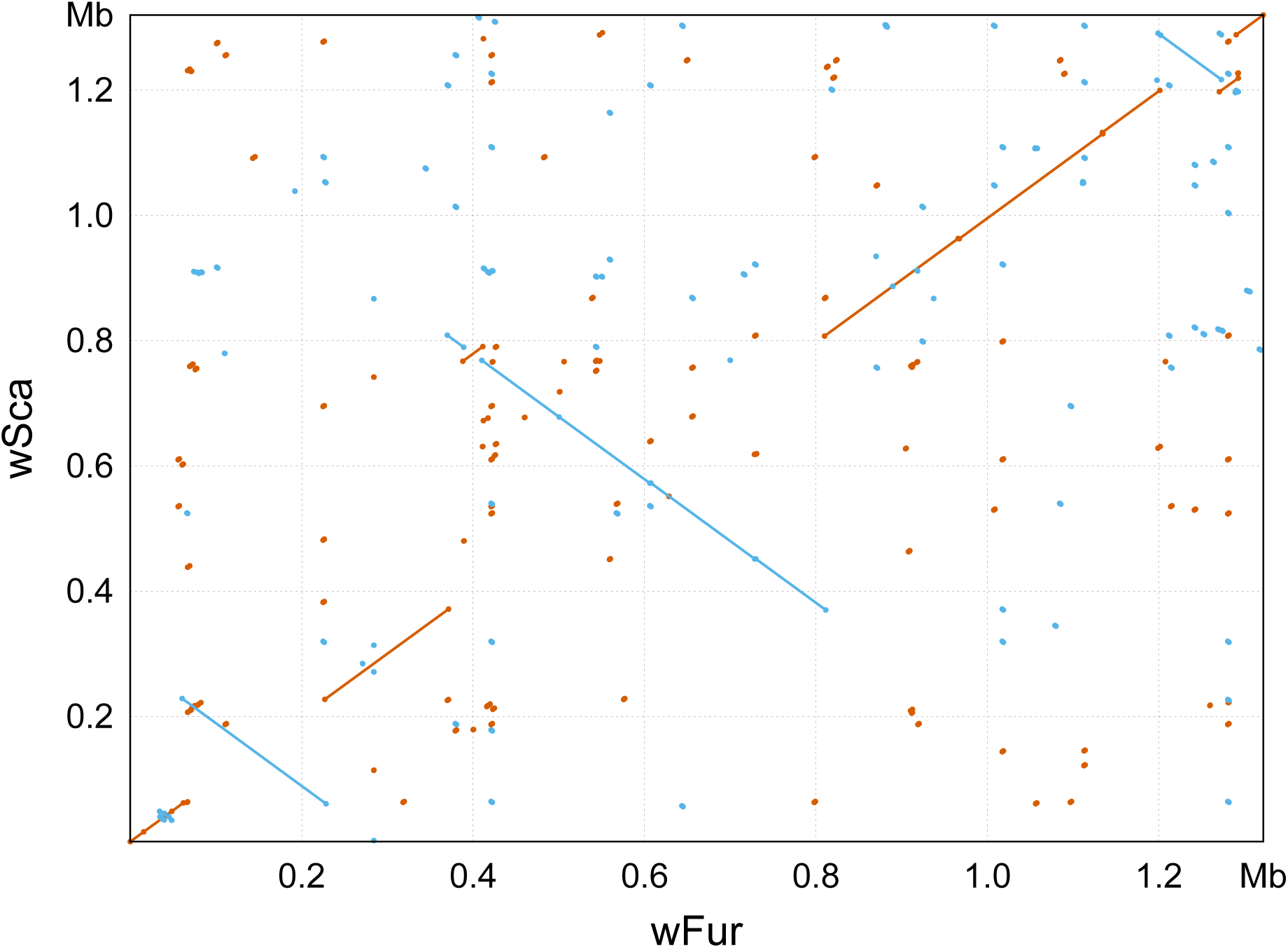
Synteny conservation between the *w*Fur and *w*Sca genomes. Dots and lines represent the alignments generated by nucmer program. Forward matches are shown in red, while reverse matches are shown in blue.

### 5. Analysis of the mitochondrial genome

The genome assemblies included information about both the hosts and *Wolbachia*. Mitochondria, in particular, are maternally inherited elements that are descended together with *Wolbachia* in infected lineages. Therefore, we analyzed mitochondrial genomes to gain insight into an evolutionary perspective of the host and symbiont. The mitochondrial genome of *w*Fur- infected *O. furnacalis* was successfully extracted from the assembly using blastn, and constructed as a single-coverage complete contig. In contrast, since the mitochondrial genome of *O. scapulalis* was not found in the Canu assembly, it was reconstructed by mapping the short-reads to the mitochondrial genome of *w*Fur-infected *O. furnacalis*. The resultant mitochondrial genomes are 15,266 bp for *w*Fur-infected *O. furnacalis* and 15,249 bp for *w*Sca-infected *O. scapulalis*.

The ANI between these two mitogenomes was 99.90%, which was greater than the ANI between normal *O. furnacalis* and *O. scapulalis* (98.77%–98.78%) (Fig. 5). Between the mitogenomes of *w*Fur-infected *O. furnacalis* and *w*Sca-infected *O. scapulalis*, we detected 14 SNPs (all of which were transitions; no transversions were detected) and several indels (1 base deletion, 2 base insertion, and 18 base deletion in the *O. scapulalis* mitochondrial genome compared to that of *O. furnacalis*, of which the last two were in AT-rich region). At the protein level, we found only two amino acid substitutions in the entire mitochondrial protein sequence. On the other hand, the mitochondrial haplotypes of infected *Ostrinia* differed substantially from those of uninfected counterparts. For example, 45 substitutions in whole protein sequences deduced from the mitochondrial genomes exist between normal (Genbank accession: MN793323) and *w*Fur-infected *O. furnacalis*. These findings indicate that *Wolbachia*-infected *Ostrinia* species have a distinct mitochondrial haplotype from uninfected individuals, which likely reflects a long period of maternal lineage separation.

**Fig. 5.**
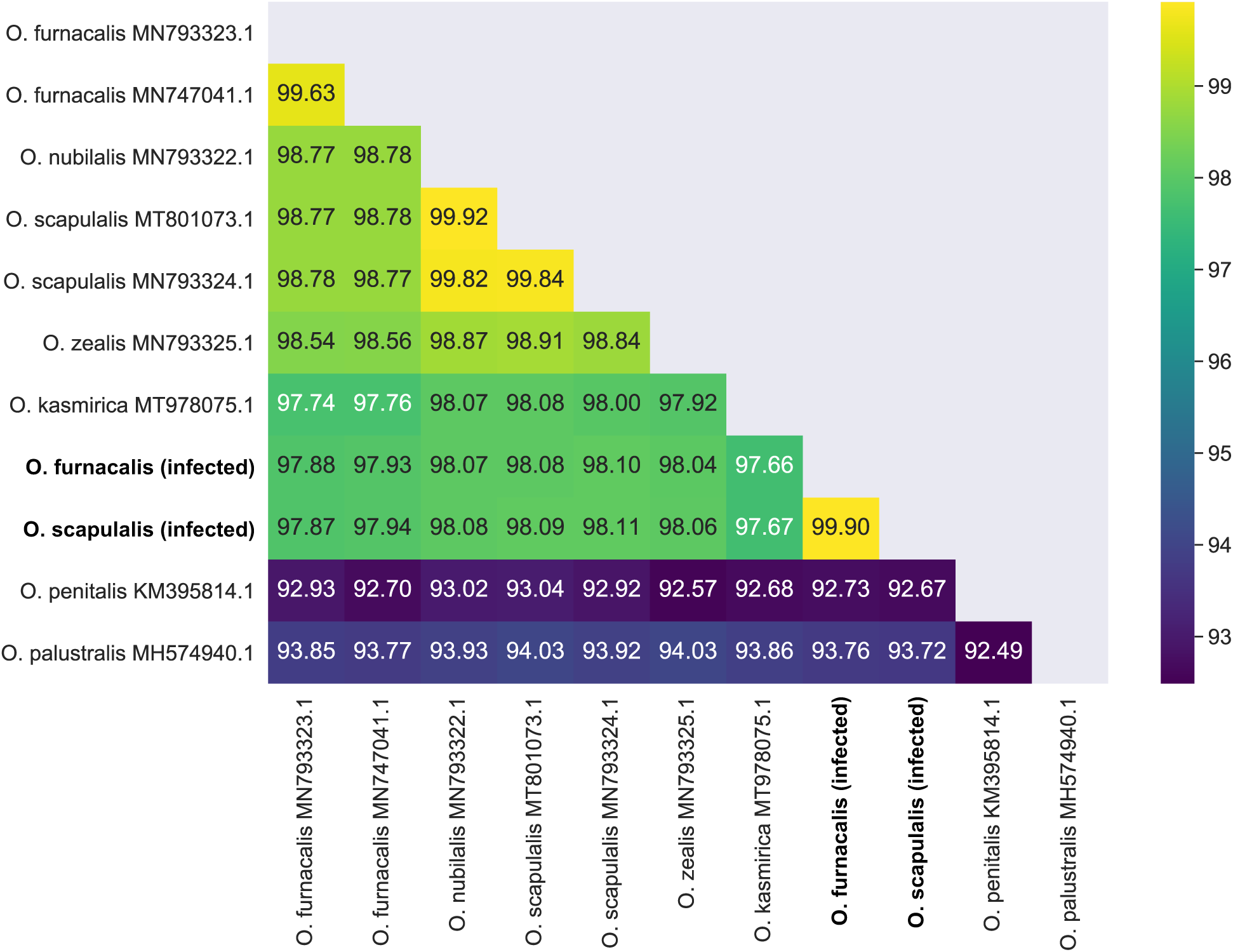
Graphical representation of ANI values for mitochondrial genomes of *Ostrinia* species. ANI values between 11 mitochondrial genomes of *Ostrinia* moths are calculated by fastANI. The sequence determined in this study are indicated in bold.

Phylogenetic analysis was performed using mitochondrial genomes of related species obtained from NCBI Genbank. Both infected lineages formed a cluster, which was located outside Clade III *Ostrinia* species including *O. furnacalis*, *O. scapulalis*, *O. nubilalis*, *O. zealis* and *O. kasmirica* (Fig. 6). We conducted a phylogenetic analysis using either nucleotide or protein sequences, and found that the topology of resultant ML trees was identical except for the phylogenetic relationship between infected lineages and *O. kasmirica* and the detailed intraclade relationship within Clade III *Ostrinia* species (Supplementary Fig. S4). Essentially, the inferred trees were consistent with previous reports (Gschloessl et al. 2020; Luo et al. 2021). According to these results, we conclude that the mitochondrial haplotype associated with *Wolbachia* infection originated in an ancestral population of Clade III *Ostrinia* species (probably excluding *O. kasmirica* based on phylogeny estimated by nucleotide sequences; Supplementary Fig. S4). Given that *Wolbachia* and mitochondria are maternally related, we can infer that *Wolbachia* infection was established in the ancestral population prior to the speciation in Clade III *Ostrinia*.

**Fig. 6.**
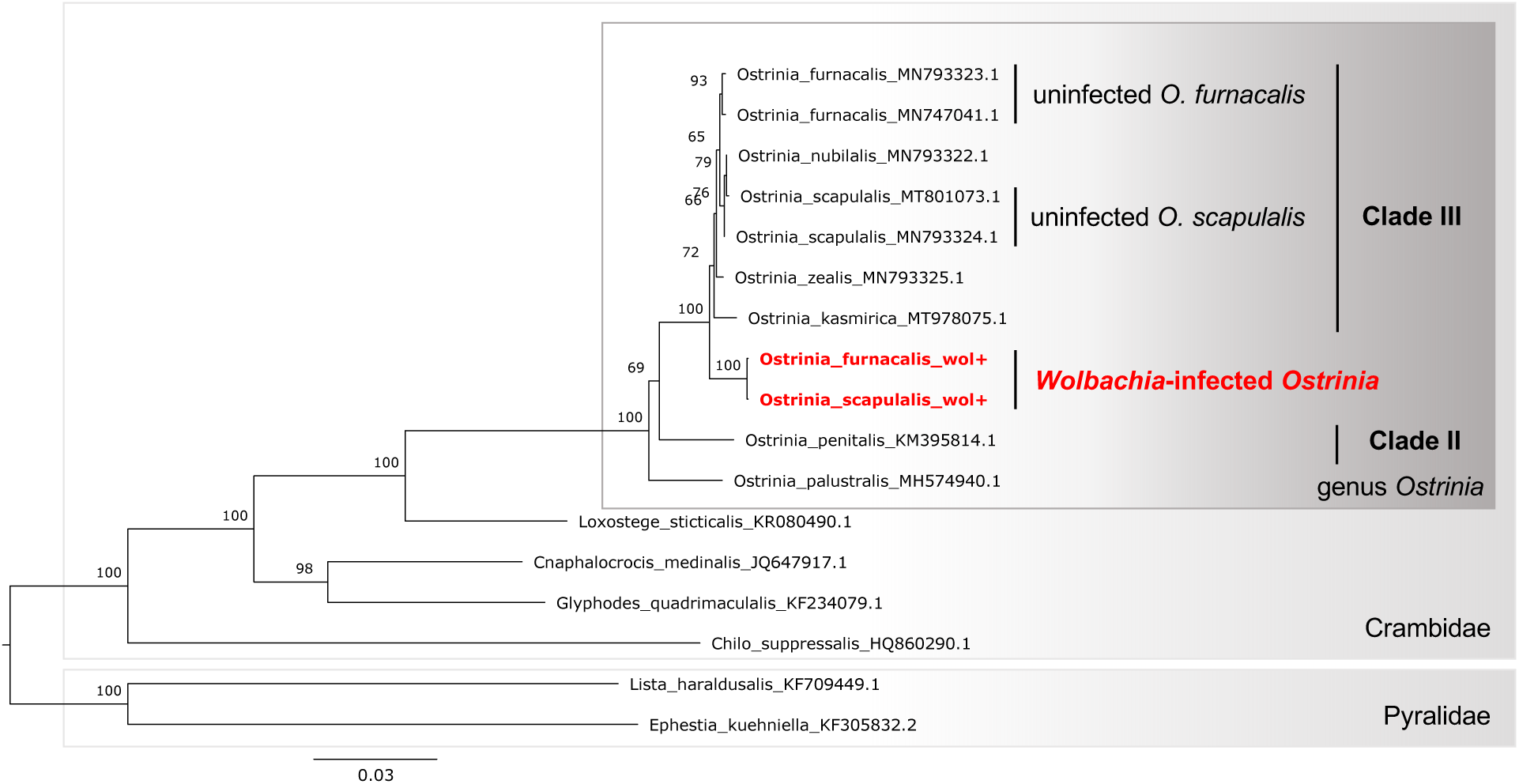
Phylogenetic relationship of mitochondrial genomes of *Ostrinia* and allied moth species. The maximum likelihood tree constructed from concatenated sequences of 13 proteins with partitions for each protein is shown. Two pyralid species (*Lista haraldusalis* and *Ephestia kuehniella*) were used as outgroups. Branch support calculated using 1000 replicates of ultrafast bootstrap is shown on the nodes. The sequence determined in this study are highlighted in red and bold.

## Discussion

We successfully sequenced the genomes of two closely related *Wolbachia* strains, *w*Fur and *w*Sca, which represent the first complete genomes of male-killing *Wolbachia* in lepidopteran hosts. The two genomes shared remarkably high homology, with over 95% of protein sequences identical. On the other hand, there are substantial differences between the genomes, most notably some large inversions (Fig. 4). It is well established that *Wolbachia* genomes poorly retain synteny between distant strains, particularly parasitic strains found in arthropod hosts (Comandatore et al. 2015; Newton et al. 2016). We observed nearly minimal transition of genome structure in two male-killing *Wolbachia* from *Ostrinia*.

The *w*Fur and *w*Sca genomes encode at least 29 and 32 ankyrin repeat-containing proteins, respectively (confirmed by hmmscan). The ankyrin repeat motif is a 33 amino acid sequence that plays a role in protein-protein interaction. Although ankyrin repeat-containing proteins are found predominantly in eukaryotes, they are also used as effectors by various pathogenic and symbiotic microbes (Al-Khodor et al. 2010). Certain genes encoding ankyrin repeat-containing proteins are found only in one of the two strains, implying that ankyrin repeat-containing proteins evolved rapidly in *Wolbachia*. The features identified by comparing closely related *Wolbachia* strains (*w*Fur and *w*Sca), namely rampant genome structure rearrangement and rapid evolution of ankyrin repeat-containing proteins, most likely represent a trend of minute genome evolution in *Wolbachia*.

The phylogenetic analysis identified *Wolbachia* of *S. picta* (the lily caterpillar) as the closest relative of *w*Fur and *w*Sca (Fig. 3). This implies the presence of *Wolbachia* host shifts between these lepidopteran hosts in evolutionary time scale. The second most closely related strain was *w*Tpre, a *Wolbachia* strain found in the parasitoid wasp *T. pretiosum*. Given the close ecological relationship between parasitic wasps and their hosts, including moths, the high sequence homology detected among *Wolbachia* species possibly indicate *Wolbachia* transmission among lepidopteran insects and their parasites. Some studies showed that horizontal transmission of *Wolbachia* is possible with the involvement of parasitic wasps (Ahmed, Li, et al. 2015; Huigens et al. 2000). Thus, wasps may be a major driver of macro-scale dynamics of *Wolbachia*, even though interspecies transmission is less well understood.

Mitochondrial analysis of *Wolbachia*-infected lineages revealed an evolutionary history of symbiotic association in the *Ostrinia* clade. Because mitochondria are inherited maternally, infected lineages’ mitochondrial haplotypes inevitably descended with *Wolbachia*. Thus, the mitochondrial phylogeny of infected lineages provides information about the time of infection establishment, from which the association of a particular mitochondrial haplotype with *Wolbachia* began. Particularly in *Ostrinia*, the removal of *Wolbachia* results in female-specific death (Kageyama & Traut 2004; also confirmed in this study), which limits the emergence of *Wolbachia*-free matrilines that retain the original *Wolbachia*-associated mitochondrial haplotype. This unique feature likely underlies the distinctiveness of mitochondrial phylogeny in infected *Ostrinia* moths. Therefore, as a result of the phylogenetic tree, we can infer that the infection was established before the speciation of Clade III *Ostrinia*, which includes *O. furnacalis* and *O. scapulalis*.

The ANI between mitochondrial haplotypes of *w*Fur-infected *O. furnacalis* and *w*Sca-infected *O. scapulalis* was greater than that between uninfected counterparts (Fig. 5). This demonstrates that *Wolbachia*-infected *O. furnacalis* and *O. scapulalis* have unusually similar mitochondrial haplotypes compared to their divergence time. Among the possible modes of *Wolbachia* dynamics (Cooper et al. 2019; Raychoudhury et al. 2009), the situation is best explained by the introgressive transmission of *Wolbachia*. The cross between *O. furnacalis* and *O. scapulalis* produces fertile hybrids that exhibit both parents’ sex pheromone types in a laboratory setting (Sakai et al. 2009). Besides, in natural populations in China, presumed hybrid individuals and introgressed genotypes have been detected (Bourguet et al. 2014). These reports demonstrate the incompleteness of reproductive isolation between these two species, which supports the possibility of *Wolbachia* introgression.

We now propose an evolutionary history model of the symbiotic association between *Ostrinia* species and *Wolbachia* based on the preceding discussion (Fig. 7): (i) *Wolbachia* infection was established in the ancestral population prior to Clade III *Ostrinia* speciation. (ii) Subsequently, Clade III *Ostrinia* species such as *O. furnacalis*, *O. scapulalis*, and others diversified, and at least one species retained *Wolbachia* infection. (iii) *Wolbachia* was recently transmitted via introgression between *O. furnacalis* and *O. scapulalis*.

**Fig. 7.**
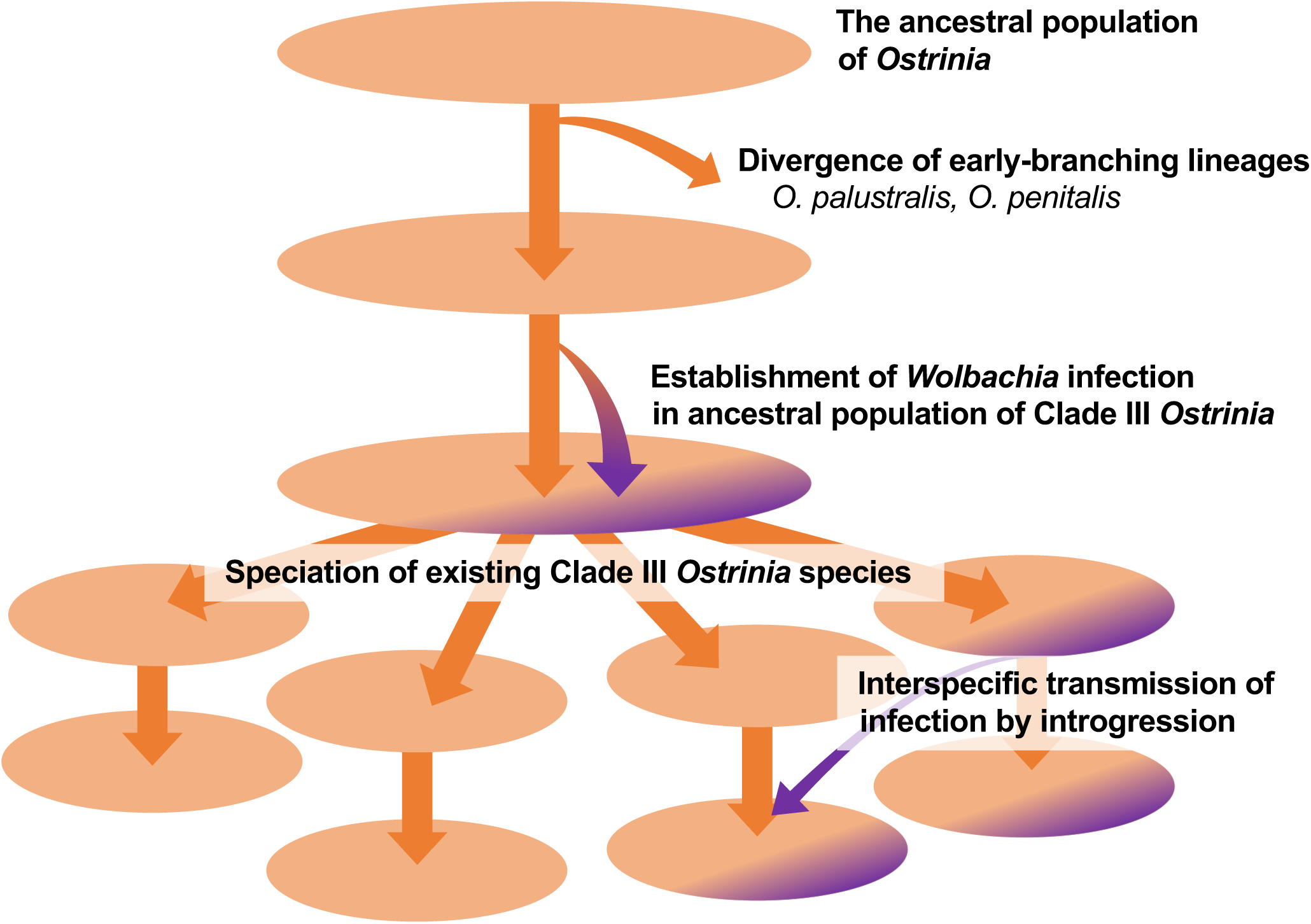
A proposed model for evolutionary history of *Wolbachia* infection in *Ostrinia* moths. Each circle represents an assumed individual species in evolutionary time scale. *Wolbachia*- infected subpopulations are depicted in a dark color.

While the outline of an evolutionary scenario was inferred, there are still unanswered questions that should be addressed in future work. First, what is the real origin of *Wolbachia* in *Ostrinia* moths? The phylogenetic tree clearly depicts the evolutionary relationship between *Wolbachia* associated with *Ostrinia* and other moth- and parasitoid wasp-associated *Wolbachia*, implying the possibility of interspecies transfer. However, since *Wolbachia* was estimated to be present in roughly half of all terrestrial arthropods (Weinert et al. 2015), the current analysis contained only a fraction of *Wolbachia*’s total diversity. A thorough survey of *Wolbachia* diversity within the Lepidoptera and wasp clades, as well as possibly other clades, is required to characterize interspecies dynamics of *Wolbachia*. *Ostrinia*-associated strains with a different origin than *w*Fur and *w*Sca may provide insight into interspecies dynamics. For instance, *Wolbachia* strains belonging to Supergroup A have been reported from *O. furnacalis* populations in China (Li et al. 2013).

Second, we sought to determine the direction of *O. furnacalis* and *O. scapulalis* introgression. Unidirectional mitochondrial introgression from *O. furnacalis* into *O. scapulalis* has been observed in Chinese populations (Bourguet et al. 2014). This indicates that the direction’s introgressive *Wolbachia* transmission was more likely. Given the high degree of homology between *w*Fur and *w*Sca or the hosts’ mitogenomes, and the resulting assumption of recent introgression, such transmission and hybridization may frequently occur among *Ostrinia* species. Indeed, it is unclear whether infected *Ostrinia* populations are genetically isolated clearly along with classical species such as *O. furnacalis* and *O. scapulalis* or they are spread across multiple species with incomplete reproductive isolation.

In conclusion, we determined two complete genomes of male-killing *Wolbachia* in two closely related lepidopteran hosts. These sequences were used to characterize nearly minimal genome evolution of *Wolbachia*. The mitochondrial genome analysis facilitated our investigation of the evolutionary relationship between *Ostrinia* moths and *Wolbachia*. It revealed *Wolbachia*’s complex dynamics across multiple hosts, including the infection establishment before allied species diversification and the introgressive transmission following host speciation. The genomic data on male-killing *Wolbachia* obtained in this study will serve as a foundation for future research on host-symbiont interaction.

## Acknowledgements

We thank the Institute for Sustainable Agro-ecosystem Services, The University of Tokyo, for collecting *Ostrinia* moths. Computations were partially performed on the NIG supercomputer at ROIS National Institute of Genetics. This work was supported by Grants-in-Aid for Scientific Research on Innovative Areas “Spectrum of the Sex: a continuity of phenotypes between female and male” (17H06431) to S.K. and T.K., Grants-in-Aid for Scientific Research (A) (22H00366) to S.K., and U TOKYO Sustainable Agriculture Education Program to T.M.

## Author contributions

T.M., T.K., and S.K. designed and performed the experiments. T.M. and H.H. performed bioinformatics analysis. T.F. performed sex pheromone analysis. T.M. and S.K. wrote the manuscript with intellectual input from all authors. S.K. supervised the project.

## Data availability

All data are available in the main text or the supplemental materials. The genome sequences of *w*Fur and *w*Sca have been deposited in GenBank under accession numbers CP096925 and CP096926, respectively.

---

**Fig. S1.**
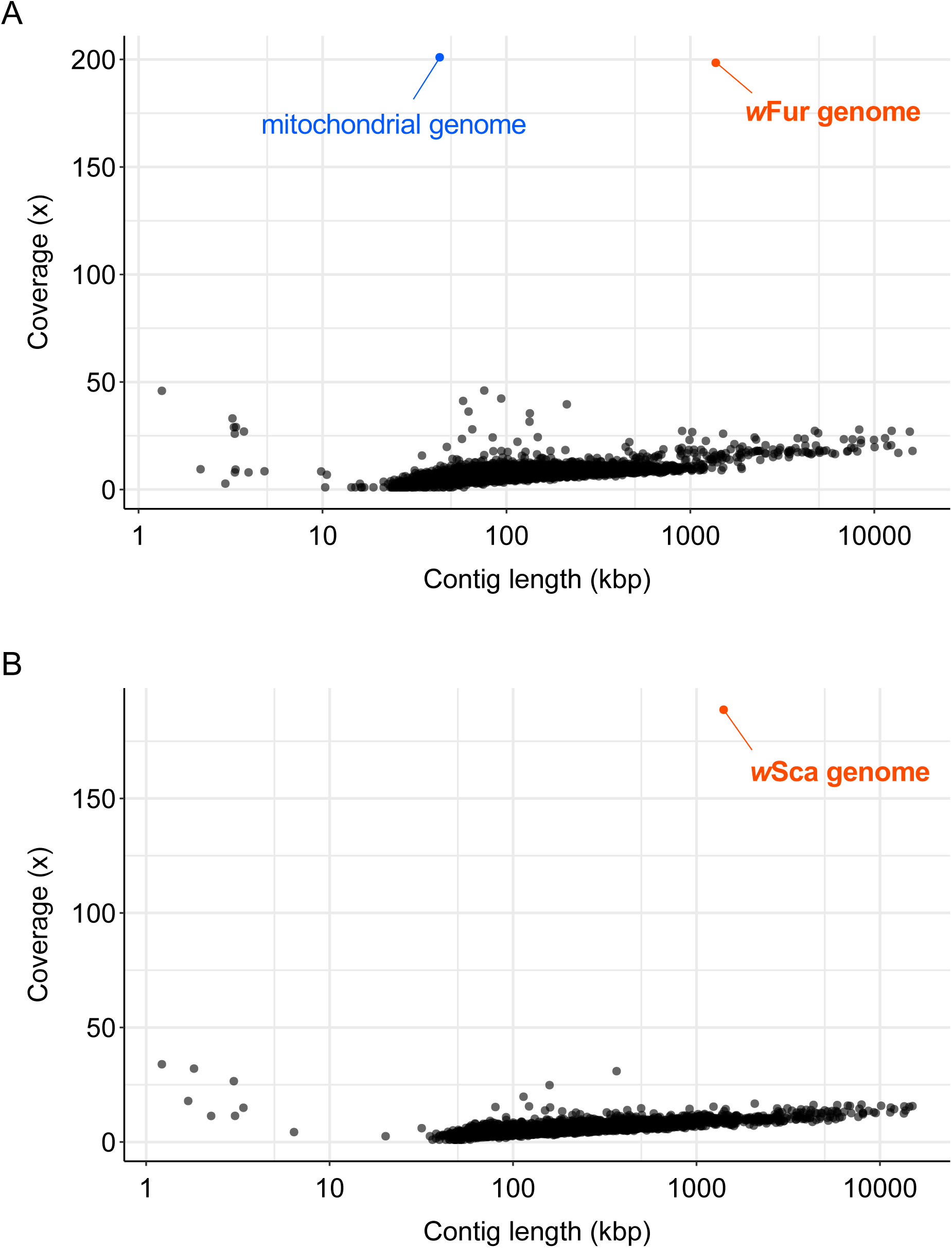
Coverage distribution of long-read assembly. Dots represent individual contigs in the genome assembly for (A) *w*Fur-infected *O. furnacalis* and (B) *w*Sca-infected *O. scapulalis* constructed by Canu. Note that a contig corresponding to the mitochondrial genome (shown in blue) is approximately 2.8 times longer than the actual mitochondrial genome due to repetitive assembly of the genome.

**Fig. S2.**
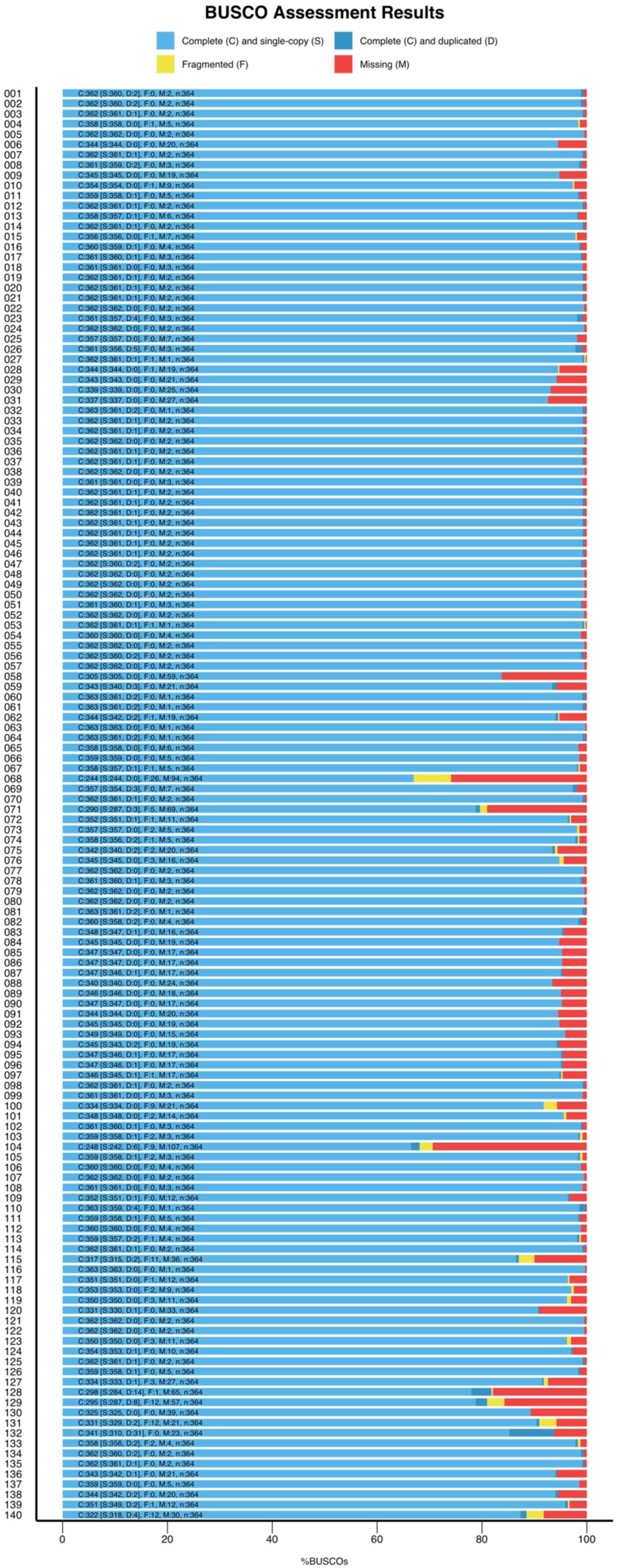
Quality evaluation of *Wolbachia* genome assemblies. A graphical representation of the BUSCO analysis in protein mode for 140 assemblies is shown. Each assembly is labeled with the identification number found in supplementary table S1.

**Fig. S3.**
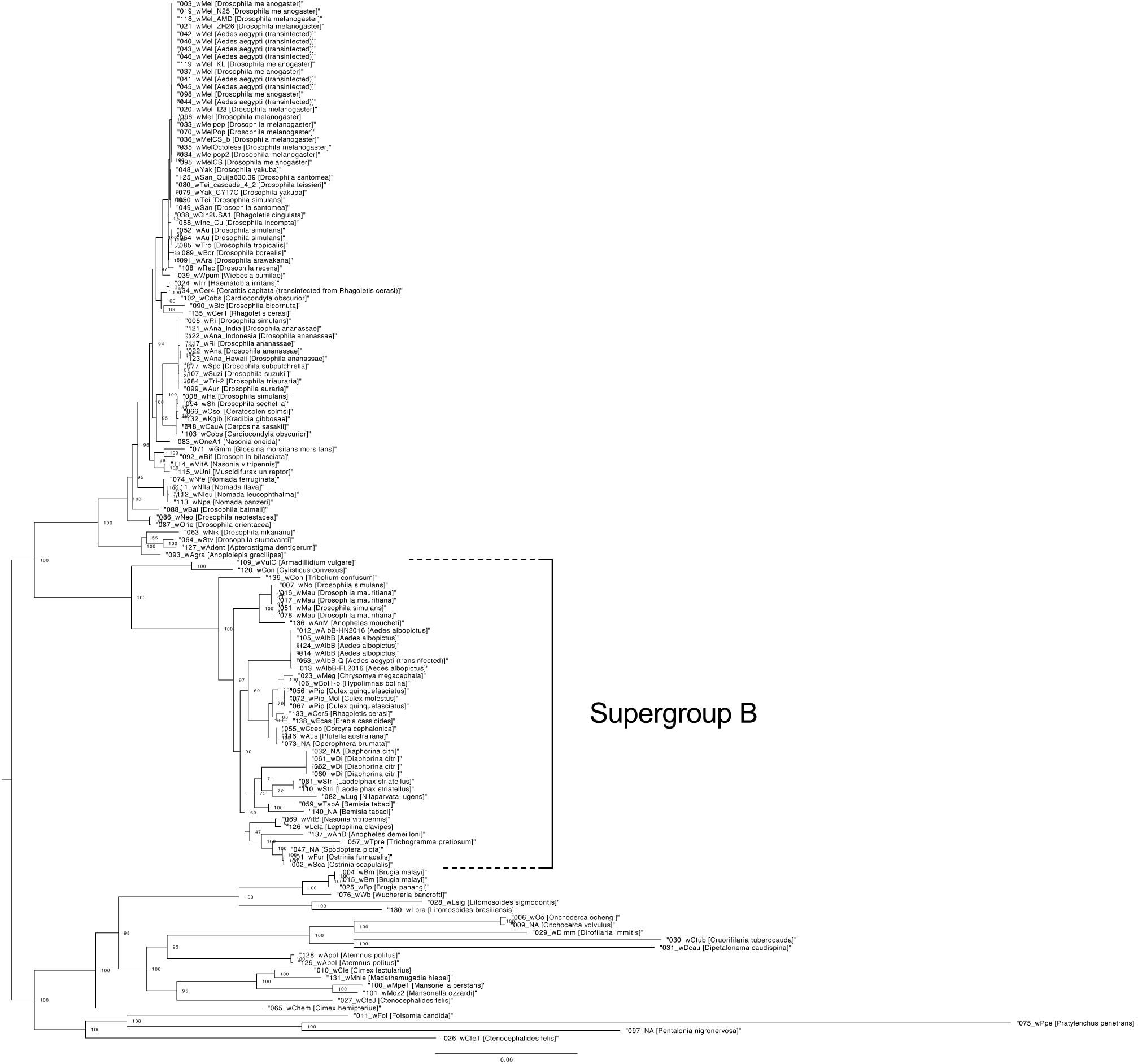
Phylogenetic relationship of 138 *Wolbachia* genomes. The maximum likelihood tree constructed from concatenated protein sequences of 63 single-copy orthologs is shown. The identification numbers, strain names, and host species are labeled. If no suitable strain name is available, it is denoted by “NA”. Branch support calculated using 1000 replicates of ultrafast bootstrap is shown on the nodes.

**Fig. S4.**
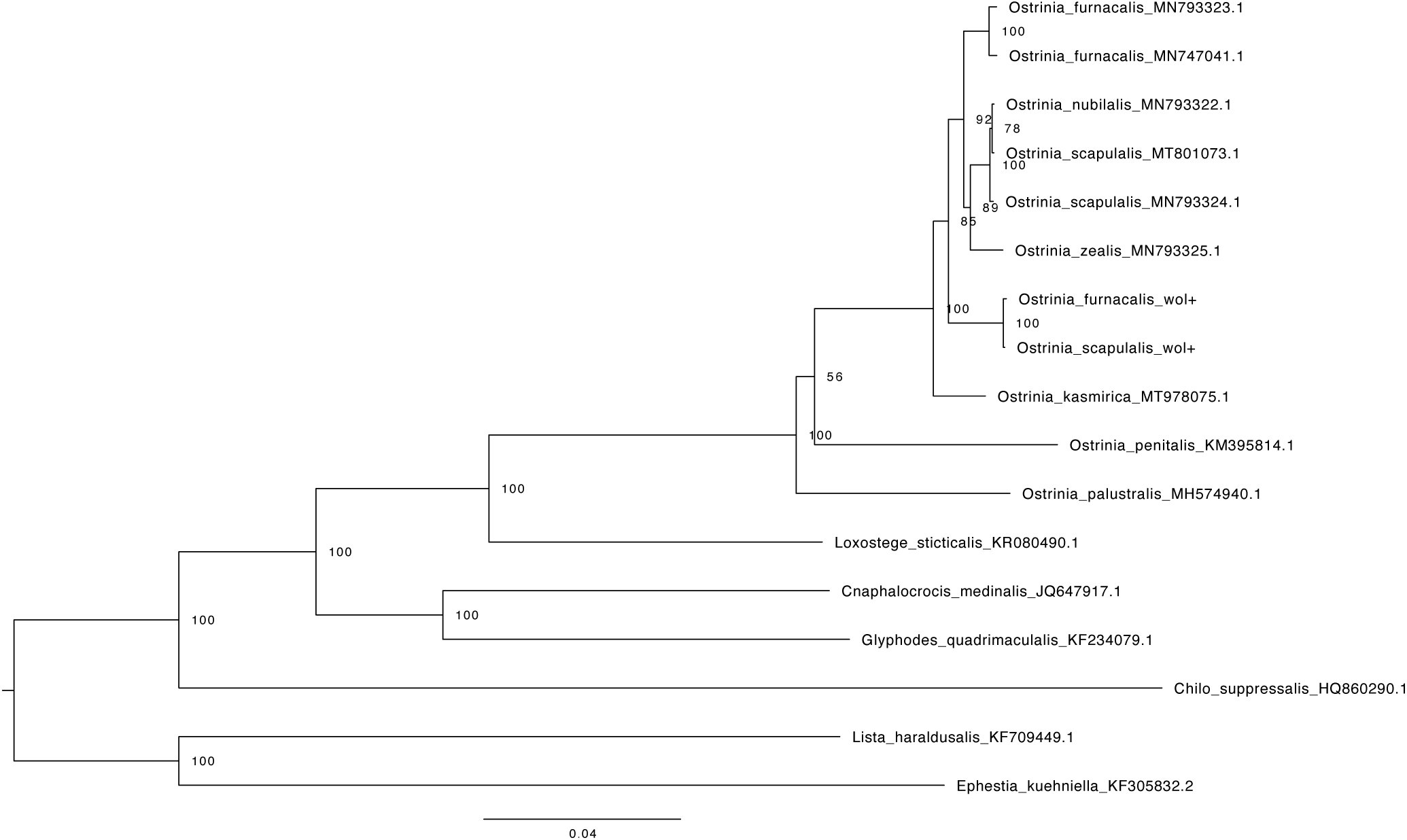
Phylogenetic relationship of mitochondrial genomes of *Ostrinia* and allied moth species based on nucleotide sequences. The maximum likelihood tree constructed from concatenated nucleotide sequences of 13 protein-coding genes, tRNAs and rRNAs is shown. Two pyralid species (*Lista haraldusalis* and *Ephestia kuehniella*) were used as outgroups. Branch support calculated using 1000 replicates of ultrafast bootstrap is shown on the nodes.

**Supplementary table S1.**
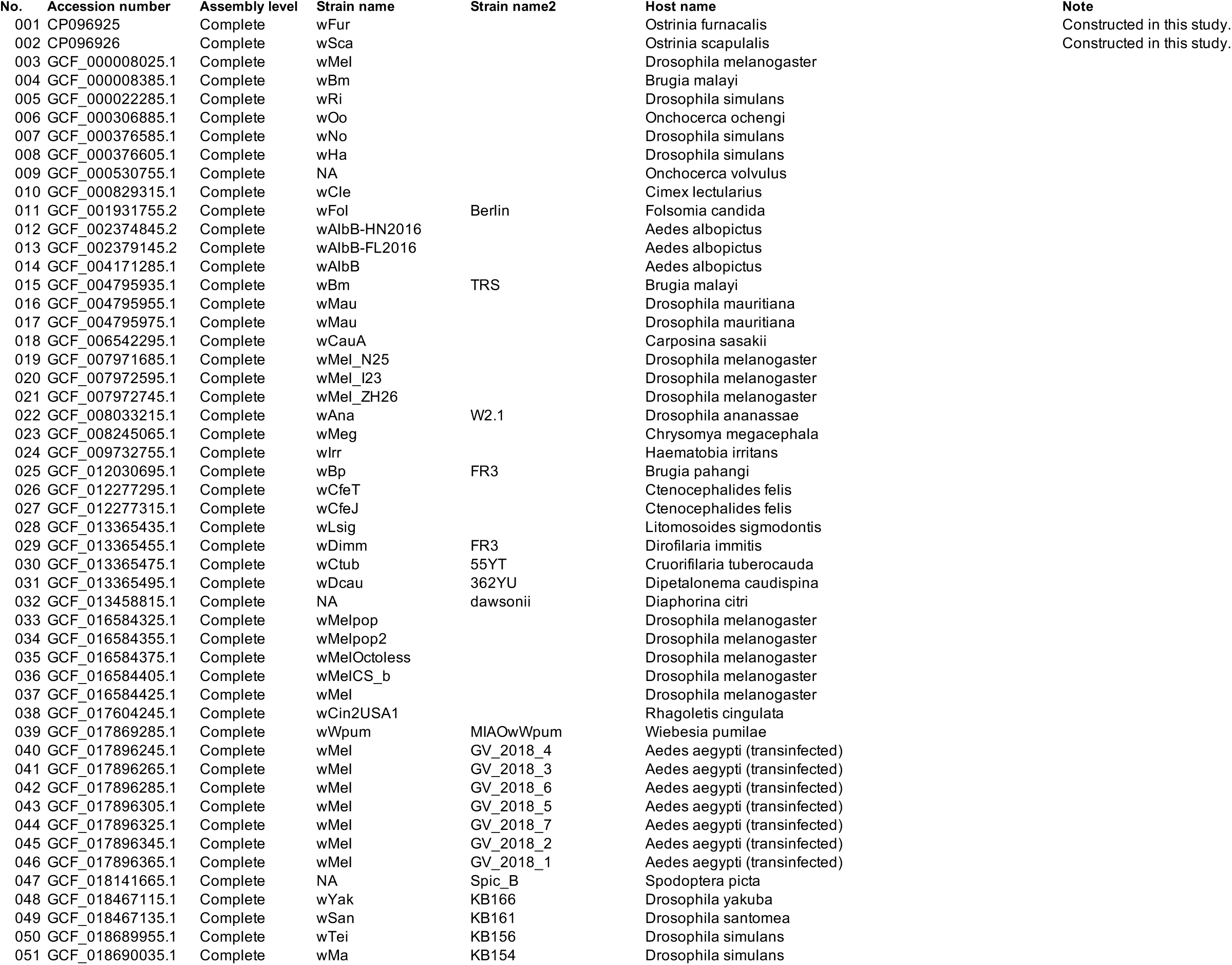
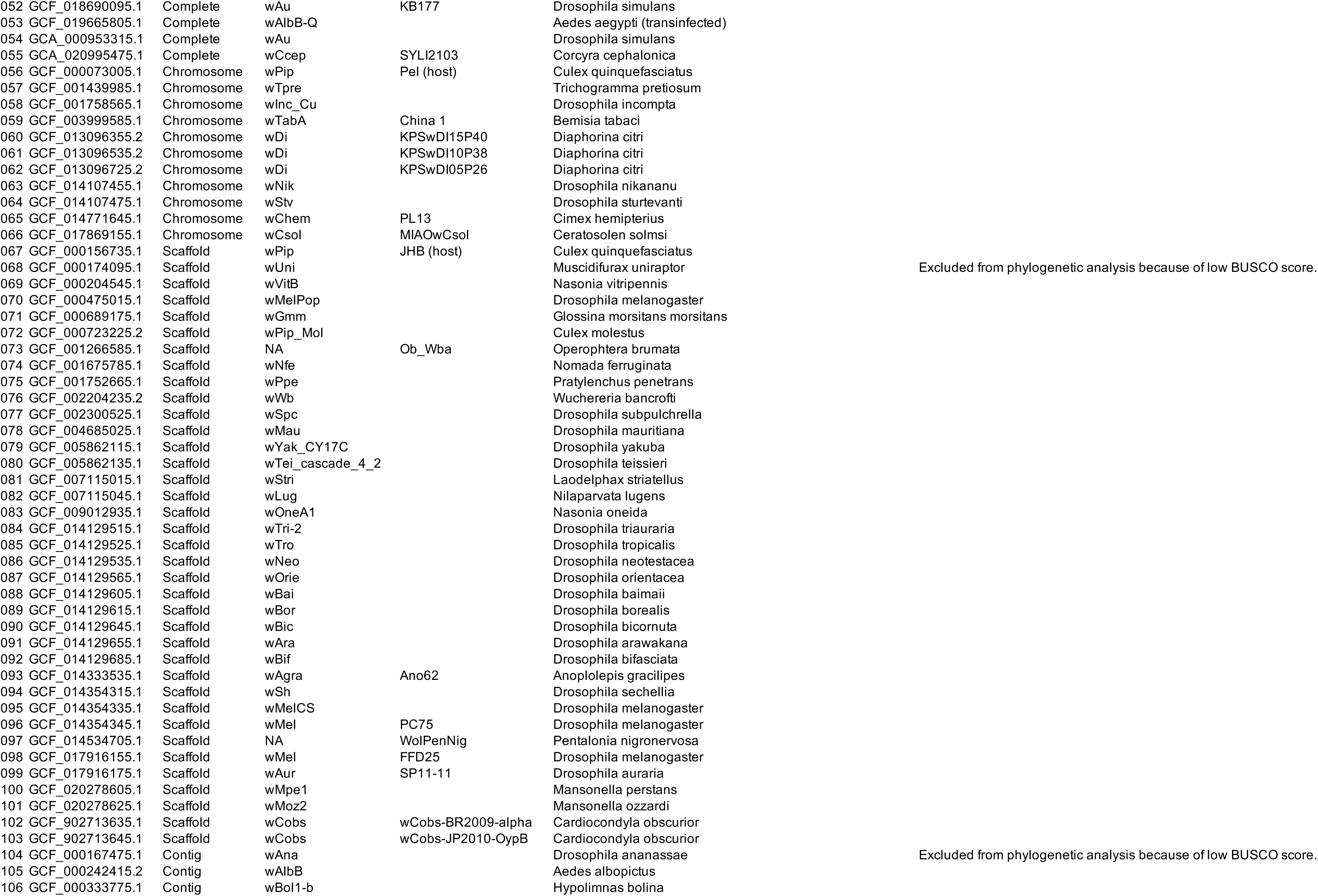
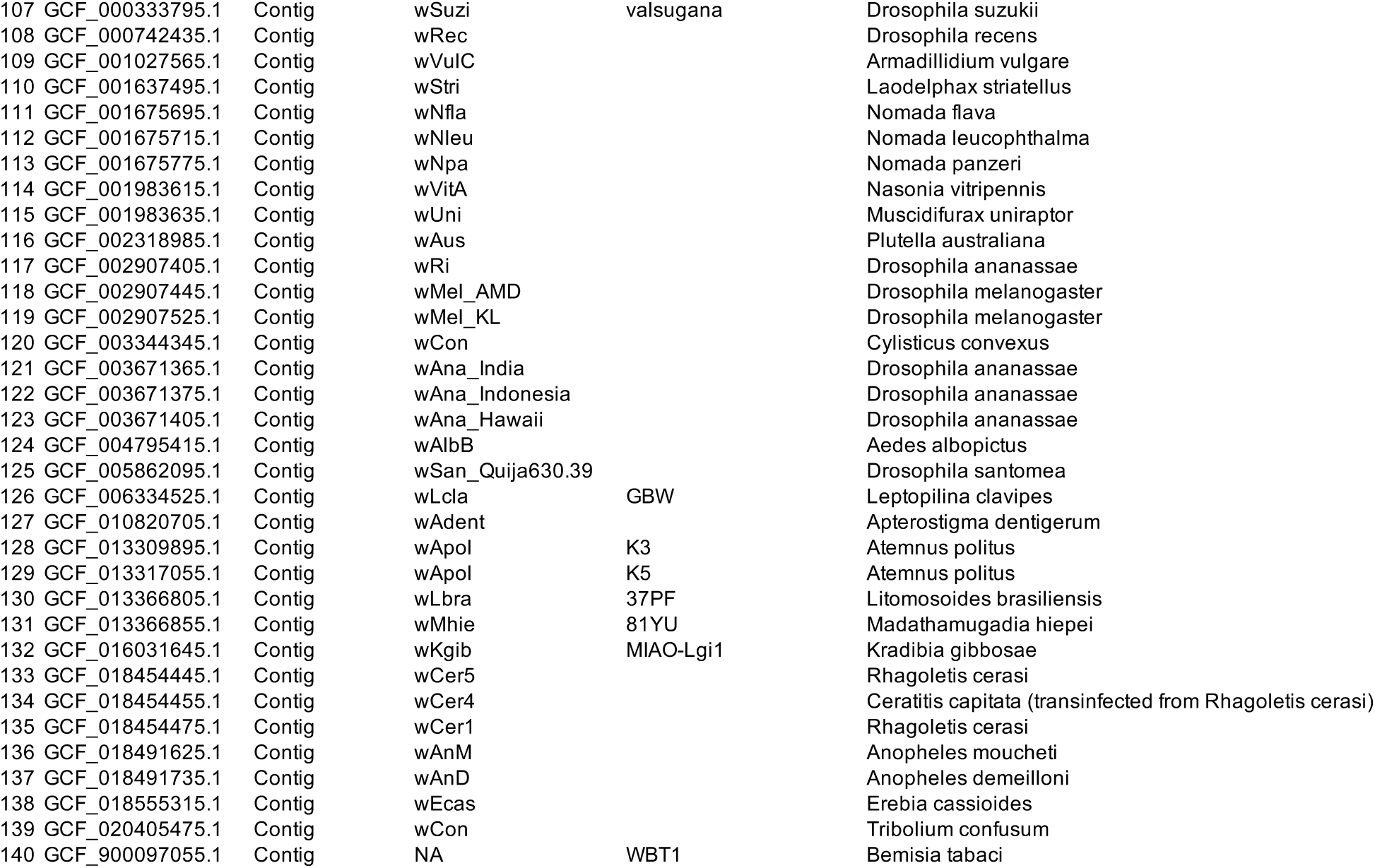
**The list of *Wolbachia* genome assemblies used in this study.** The column “Strain name2” indicates special notes on strain names of *Wolbachia* provided in each Genbank entry.

**Supplementary table S2.**
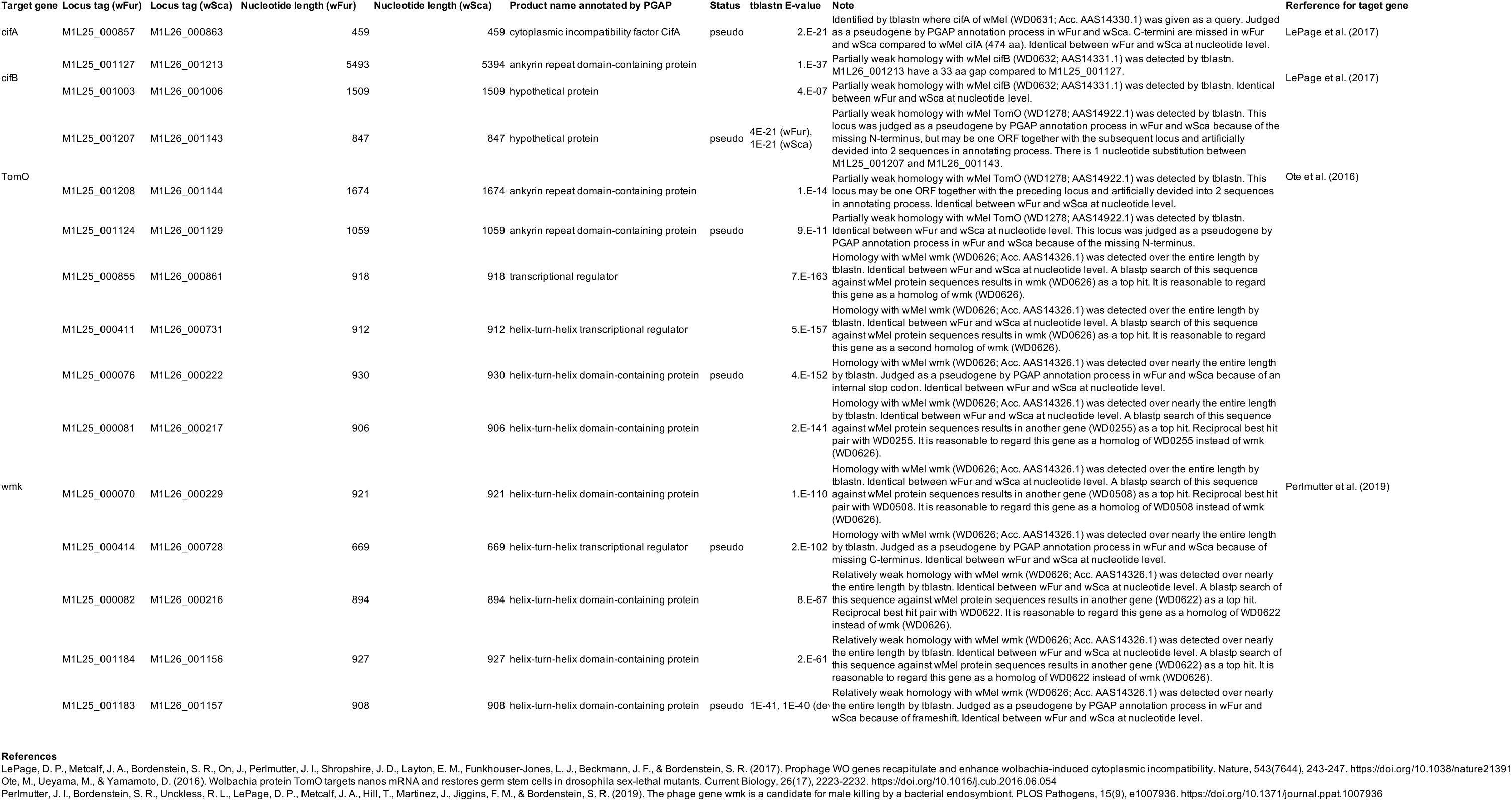
**The list of loci which exhibit homology with host-manipulating *Wolbachia* genes in *w* Fur and *w* Sca genomes.** Gene loci with E value < 1E-05 (except for wmk) in the results of tblastn search are shown. For wmk, a threshold is set as 1E-40 due to many fragmented hits. The columns “nt length” show the length of the loci (not the alignment length of tblastn).

**Supplementary table S3.**
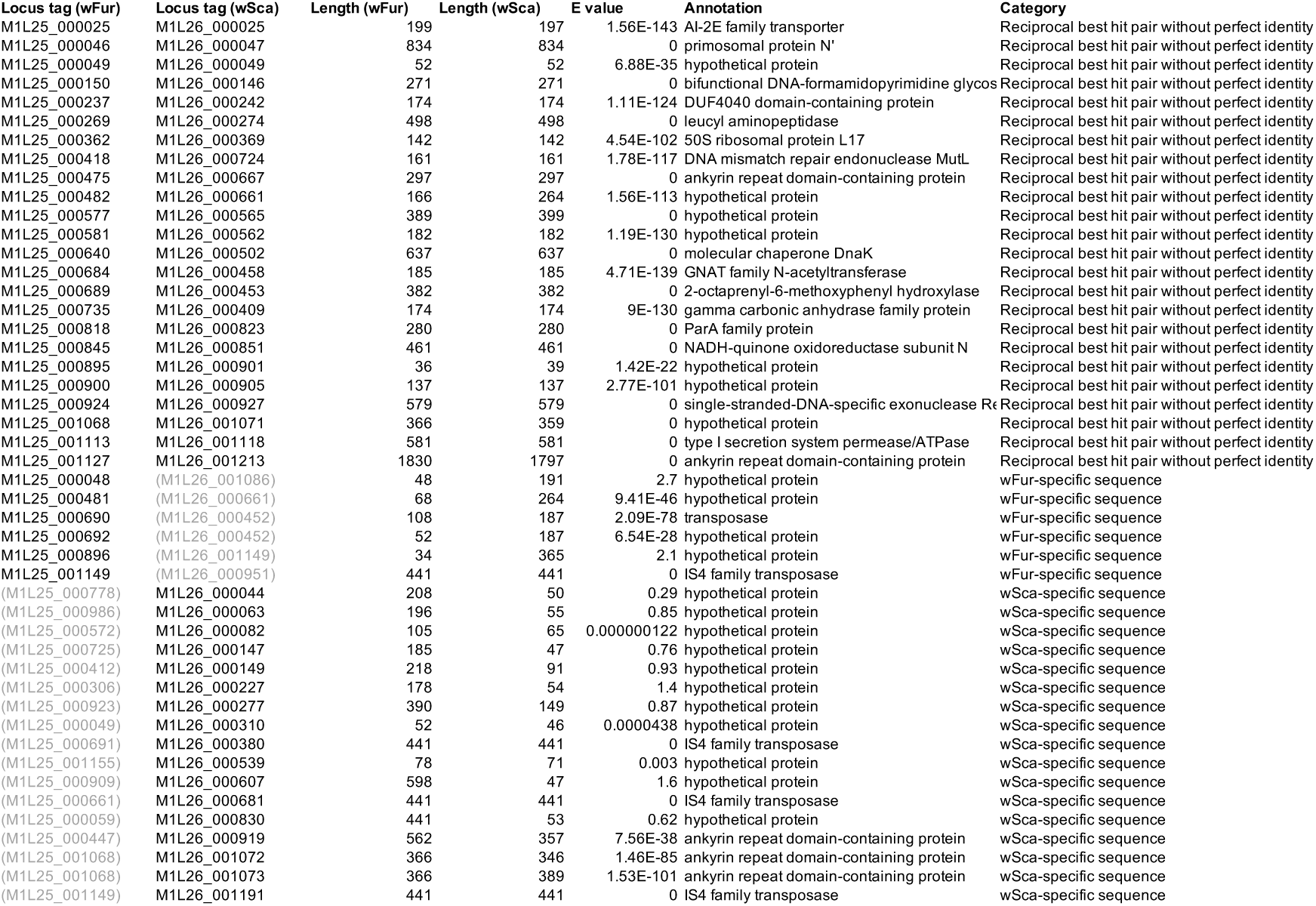
**The list of protein sequences which do not have identical counterparts between *w* Fur and *w* Sca.** For one strain-specific sequences, the subject sequences of BLASTp searc are described. We assigned “low similarity in the other’s genome” when E value is bigger than 1E-10.

## Notes

### Competing Interest Statement

The authors have declared no competing interest.

