## Supplementary material for "Two complete genomes of male-killing *Wolbachia* infecting *Ostrinia* moth species illuminate their evolutionary dynamics and association with hosts": Table S3

**Supplementary table S3. The list of protein sequences which do not have identical counterparts between *w*Fur and *w*Sca.** For one strain-specific sequences, the subject sequences of BLASTp searc are described. We assigned "low similarity in the other's genome" when E value is bigger than 1E-10.

| Locus tag (wFur) | Locus tag (wSca) | Length (wFur) | Length (wSca) | E value | Annotation | Category |
|---|---|---|---|---|---|---|
| M1L25_000025 | M1L26_000025 | 199 | 197 | 1.56E-143 | AI-2E family transporter | Reciprocal best hit pair without perfect identity |
| M1L25_000046 | M1L26_000047 | 834 | 834 | 0 | primosomal protein N' | Reciprocal best hit pair without perfect identity |
| M1L25_000049 | M1L26_000049 | 52 | 52 | 6.88E-35 | hypothetical protein | Reciprocal best hit pair without perfect identity |
| M1L25_000150 | M1L26_000146 | 271 | 271 | 0 | bifunctional DNA-formamidopyrimidine glycos | Reciprocal best hit pair without perfect identity |
| M1L25_000237 | M1L26_000242 | 174 | 174 | 1.11E-124 | DUF4040 domain-containing protein | Reciprocal best hit pair without perfect identity |
| M1L25_000269 | M1L26_000274 | 498 | 498 | 0 | leucyl aminopeptidase | Reciprocal best hit pair without perfect identity |
| M1L25_000362 | M1L26_000369 | 142 | 142 | 4.54E-102 | 50S ribosomal protein L17 | Reciprocal best hit pair without perfect identity |
| M1L25_000418 | M1L26_000724 | 161 | 161 | 1.78E-117 | DNA mismatch repair endonuclease MutL | Reciprocal best hit pair without perfect identity |
| M1L25_000475 | M1L26_000667 | 297 | 297 | 0 | ankyrin repeat domain-containing protein | Reciprocal best hit pair without perfect identity |
| M1L25_000482 | M1L26_000661 | 166 | 264 | 1.56E-113 | hypothetical protein | Reciprocal best hit pair without perfect identity |
| M1L25_000577 | M1L26_000565 | 389 | 399 | 0 | hypothetical protein | Reciprocal best hit pair without perfect identity |
| M1L25_000581 | M1L26_000562 | 182 | 182 | 1.19E-130 | hypothetical protein | Reciprocal best hit pair without perfect identity |
| M1L25_000640 | M1L26_000502 | 637 | 637 | 0 | molecular chaperone DnaK | Reciprocal best hit pair without perfect identity |
| M1L25_000684 | M1L26_000458 | 185 | 185 | 4.71E-139 | GNAT family N-acetyltransferase | Reciprocal best hit pair without perfect identity |
| M1L25_000689 | M1L26_000453 | 382 | 382 | 0 | 2-octaprenyl-6-methoxyphenyl hydroxylase | Reciprocal best hit pair without perfect identity |
| M1L25_000735 | M1L26_000409 | 174 | 174 | 9E-130 | gamma carbonic anhydrase family protein | Reciprocal best hit pair without perfect identity |
| M1L25_000818 | M1L26_000823 | 280 | 280 | 0 | ParA family protein | Reciprocal best hit pair without perfect identity |
| M1L25_000845 | M1L26_000851 | 461 | 461 | 0 | NADH-quinone oxidoreductase subunit N | Reciprocal best hit pair without perfect identity |
| M1L25_000895 | M1L26_000901 | 36 | 39 | 1.42E-22 | hypothetical protein | Reciprocal best hit pair without perfect identity |
| M1L25_000900 | M1L26_000905 | 137 | 137 | 2.77E-101 | hypothetical protein | Reciprocal best hit pair without perfect identity |
| M1L25_000924 | M1L26_000927 | 579 | 579 | 0 | single-stranded-DNA-specific exonuclease Re | Reciprocal best hit pair without perfect identity |
| M1L25_001068 | M1L26_001071 | 366 | 359 | 0 | hypothetical protein | Reciprocal best hit pair without perfect identity |
| M1L25_001113 | M1L26_001118 | 581 | 581 | 0 | type I secretion system permease/ATPase | Reciprocal best hit pair without perfect identity |
| M1L25_001127 | M1L26_001213 | 1830 | 1797 | 0 | ankyrin repeat domain-containing protein | Reciprocal best hit pair without perfect identity |
| M1L25_000048 | (M1L26_001086) | 48 | 191 | 2.7 | hypothetical protein | wFur-specific sequence |
| M1L25_000481 | (M1L26_000661) | 68 | 264 | 9.41E-46 | hypothetical protein | wFur-specific sequence |
| M1L25_000690 | (M1L26_000452) | 108 | 187 | 2.09E-78 | transposase | wFur-specific sequence |
| M1L25_000692 | (M1L26_000452) | 52 | 187 | 6.54E-28 | hypothetical protein | wFur-specific sequence |
| M1L25_000896 | (M1L26_001149) | 34 | 365 | 2.1 | hypothetical protein | wFur-specific sequence |
| M1L25_001149 | (M1L26_000951) | 441 | 441 | 0 | IS4 family transposase | wFur-specific sequence |
| (M1L25_000778) | M1L26_000044 | 208 | 50 | 0.29 | hypothetical protein | wSca-specific sequence |
| (M1L25_000986) | M1L26_000063 | 196 | 55 | 0.85 | hypothetical protein | wSca-specific sequence |
| (M1L25_000572) | M1L26_000082 | 105 | 65 | 0.000000122 | hypothetical protein | wSca-specific sequence |
| (M1L25_000725) | M1L26_000147 | 185 | 47 | 0.76 | hypothetical protein | wSca-specific sequence |
| (M1L25_000412) | M1L26_000149 | 218 | 91 | 0.93 | hypothetical protein | wSca-specific sequence |
| (M1L25_000306) | M1L26_000227 | 178 | 54 | 1.4 | hypothetical protein | wSca-specific sequence |
| (M1L25_000923) | M1L26_000277 | 390 | 149 | 0.87 | hypothetical protein | wSca-specific sequence |
| (M1L25_000049) | M1L26_000310 | 52 | 46 | 0.0000438 | hypothetical protein | wSca-specific sequence |
| (M1L25_000691) | M1L26_000380 | 441 | 441 | 0 | IS4 family transposase | wSca-specific sequence |
| (M1L25_001155) | M1L26_000539 | 78 | 71 | 0.003 | hypothetical protein | wSca-specific sequence |
| (M1L25_000909) | M1L26_000607 | 598 | 47 | 1.6 | hypothetical protein | wSca-specific sequence |
| (M1L25_000661) | M1L26_000681 | 441 | 441 | 0 | IS4 family transposase | wSca-specific sequence |
| (M1L25_000059) | M1L26_000830 | 441 | 53 | 0.62 | hypothetical protein | wSca-specific sequence |
| (M1L25_000447) | M1L26_000919 | 562 | 357 | 7.56E-38 | ankyrin repeat domain-containing protein | wSca-specific sequence |
| (M1L25_001068) | M1L26_001072 | 366 | 346 | 1.46E-85 | ankyrin repeat domain-containing protein | wSca-specific sequence |
| (M1L25_001068) | M1L26_001073 | 366 | 389 | 1.53E-101 | ankyrin repeat domain-containing protein | wSca-specific sequence |
| (M1L25_001149) | M1L26_001191 | 441 | 441 | 0 | IS4 family transposase | wSca-specific sequence |
