## Supplementary material for "Two complete genomes of male-killing *Wolbachia* infecting *Ostrinia* moth species illuminate their evolutionary dynamics and association with hosts": Table S2

**Supplementary table S2. The list of loci which exhibit homology with host-manipulating *Wolbachia* genes in *w*Fur and *w*Sca genomes.** Gene loci with E value < 1E-05 (except for wmk) in the results of tblastn search are shown. For wmk, a threshold is set as 1E-40 due to many fragmented hits. The columns "nt length" show the length of the loci (not the alignment length of tblastn).

| Target gene | Locus tag (wFur) | Locus tag (wSca) | Nucleotide length (wFur) | Nucleotide length (wSca) | Product name annotated by PGAP | Status | tblastn E-value | Note | Rerference for taget gene |
|---|---|---|---|---|---|---|---|---|---|
| cifA | M1L25_000857 | M1L26_000863 | 459 | 459 | cytoplasmic incompatibility factor CifA | pseudo | 2.E-21 | Identified by tblastn where cifA of wMel (WD0631; Acc. AAS14330.1) was given as a query. Judged as a pseudogene by PGAP annotation process in wFur and wSca. C-termini are missed in wFur and wSca compared to wMel cifA (474 aa). Identical between wFur and wSca at nucleotide level. | LePage et al. (2017) |
| cifB | M1L25_001127 | M1L26_001213 | 5493 | 5394 | ankyrin repeat domain-containing protein | | 1.E-37 | Partially weak homology with wMel cifB (WD0632; AAS14331.1) was detected by tblastn. M1L26_001213 have a 33 aa gap compared to M1L25_001127. | LePage et al. (2017) |
| | M1L25_001003 | M1L26_001006 | 1509 | 1509 | hypothetical protein | | 4.E-07 | Partially weak homology with wMel cifB (WD0632; AAS14331.1) was detected by tblastn. Identical between wFur and wSca at nucleotide level. | |
| TomO | M1L25_001207 | M1L26_001143 | 847 | 847 | hypothetical protein | pseudo | 4E-21 (wFur), 1E-21 (wSca) | Partially weak homology with wMel TomO (WD1278; AAS14922.1) was detected by tblastn. This locus was judged as a pseudogene by PGAP annotation process in wFur and wSca because of the missing N-terminus, but may be one ORF together with the subsequent locus and artificially devided into 2 sequences in annotating process. There is 1 nucleotide substitution between M1L25_001207 and M1L26_001143. | Ote et al. (2016) |
| | M1L25_001208 | M1L26_001144 | 1674 | 1674 | ankyrin repeat domain-containing protein | | 1.E-14 | Partially weak homology with wMel TomO (WD1278; AAS14922.1) was detected by tblastn. This locus may be one ORF together with the preceding locus and artificially devided into 2 sequences in annotating process. Identical between wFur and wSca at nucleotide level. | |
| | M1L25_001124 | M1L26_001129 | 1059 | 1059 | ankyrin repeat domain-containing protein | pseudo | 9.E-11 | Partially weak homology with wMel TomO (WD1278; AAS14922.1) was detected by tblastn. Identical between wFur and wSca at nucleotide level. This locus was judged as a pseudogene by PGAP annotation process in wFur and wSca because of the missing N-terminus. | |
| wmk | M1L25_000855 | M1L26_000861 | 918 | 918 | transcriptional regulator | | 7.E-163 | Homology with wMel wmk (WD0626; Acc. AAS14326.1) was detected over the entire length by tblastn. Identical between wFur and wSca at nucleotide level. A blastp search of this sequence against wMel protein sequences results in wmk (WD0626) as a top hit. It is reasonable to regard this gene as a homolog of wmk (WD0626). | Perlmutter et al. (2019) |
| | M1L25_000411 | M1L26_000731 | 912 | 912 | helix-turn-helix transcriptional regulator | | 5.E-157 | Homology with wMel wmk (WD0626; Acc. AAS14326.1) was detected over the entire length by tblastn. Identical between wFur and wSca at nucleotide level. A blastp search of this sequence against wMel protein sequences results in wmk (WD0626) as a top hit. It is reasonable to regard this gene as a second homolog of wmk (WD0626). | |
| | M1L25_000076 | M1L26_000222 | 930 | 930 | helix-turn-helix domain-containing protein | pseudo | 4.E-152 | Homology with wMel wmk (WD0626; Acc. AAS14326.1) was detected over nearly the entire length by tblastn. Judged as a pseudogene by PGAP annotation process in wFur and wSca because of an internal stop codon. Identical between wFur and wSca at nucleotide level. | |
| | M1L25_000081 | M1L26_000217 | 906 | 906 | helix-turn-helix domain-containing protein | | 2.E-141 | Homology with wMel wmk (WD0626; Acc. AAS14326.1) was detected over nearly the entire length by tblastn. Identical between wFur and wSca at nucleotide level. A blastp search of this sequence against wMel protein sequences results in another gene (WD0255) as a top hit. Reciprocal best hit pair with WD0255. It is reasonable to regard this gene as a homolog of WD0255 instead of wmk (WD0626). | |
| | M1L25_000070 | M1L26_000229 | 921 | 921 | helix-turn-helix domain-containing protein | | 1.E-110 | Homology with wMel wmk (WD0626; Acc. AAS14326.1) was detected over the entire length by tblastn. Identical between wFur and wSca at nucleotide level. A blastp search of this sequence against wMel protein sequences results in another gene (WD0508) as a top hit. Reciprocal best hit pair with WD0508. It is reasonable to regard this gene as a homolog of WD0508 instead of wmk (WD0626). | |
| | M1L25_000414 | M1L26_000728 | 669 | 669 | helix-turn-helix transcriptional regulator | pseudo | 2.E-102 | Homology with wMel wmk (WD0626; Acc. AAS14326.1) was detected over nearly the entire length by tblastn. Judged as a pseudogene by PGAP annotation process in wFur and wSca because of missing C-terminus. Identical between wFur and wSca at nucleotide level. | |
| | M1L25_000082 | M1L26_000216 | 894 | 894 | helix-turn-helix domain-containing protein | | 8.E-67 | Relatively weak homology with wMel wmk (WD0626; Acc. AAS14326.1) was detected over nearly the entire length by tblastn. Identical between wFur and wSca at nucleotide level. A blastp search of this sequence against wMel protein sequences results in another gene (WD0622) as a top hit. Reciprocal best hit pair with WD0622. It is reasonable to regard this gene as a homolog of WD0622 instead of wmk (WD0626). | |
| | M1L25_001184 | M1L26_001156 | 927 | 927 | helix-turn-helix domain-containing protein | | 2.E-61 | Relatively weak homology with wMel wmk (WD0626; Acc. AAS14326.1) was detected over nearly the entire length by tblastn. Identical between wFur and wSca at nucleotide level. A blastp search of this sequence against wMel protein sequences results in another gene (WD0622) as a top hit. It is reasonable to regard this gene as a homolog of WD0622 instead of wmk (WD0626). | |
| | M1L25_001183 | M1L26_001157 | 908 | 908 | helix-turn-helix domain-containing protein | pseudo | 1E-41, 1E-40 (dev | Relatively weak homology with wMel wmk (WD0626; Acc. AAS14326.1) was detected over nearly the entire length by tblastn. Judged as a pseudogene by PGAP annotation process in wFur and wSca because of frameshift. Identical between wFur and wSca at nucleotide level. | |
