## Supplementary material for "Two complete genomes of male-killing *Wolbachia* infecting *Ostrinia* moth species illuminate their evolutionary dynamics and association with hosts": Table S1

**Supplementary table S1. The list of *Wolbachia* genome assemblies used in this study.** The column "Strain name2" indicates special notes on strain names of *Wolbachia* provided in each Genbank entry.

| No. | Accession number | Assembly level | Strain name | Strain name2 | Host name | Note |
|---|---|---|---|---|---|---|
| 001 | CP096925 | Complete | wFur | | Ostrinia furnacalis | Constructed in this study. |
| 002 | CP096926 | Complete | wSca | | Ostrinia scapulalis | Constructed in this study. |
| 003 | GCF_000008025.1 | Complete | wMel | | Drosophila melanogaster | |
| 004 | GCF_000008385.1 | Complete | wBm | | Brugia malayi | |
| 005 | GCF_000022285.1 | Complete | wRi | | Drosophila simulans | |
| 006 | GCF_000306885.1 | Complete | wOo | | Onchocerca ochengi | |
| 007 | GCF_000376585.1 | Complete | wNo | | Drosophila simulans | |
| 008 | GCF_000376605.1 | Complete | wHa | | Drosophila simulans | |
| 009 | GCF_000530755.1 | Complete | NA | | Onchocerca volvulus | |
| 010 | GCF_000829315.1 | Complete | wCle | | Cimex lectularius | |
| 011 | GCF_001931755.2 | Complete | wFol | Berlin | Folsomia candida | |
| 012 | GCF_002374845.2 | Complete | wAlbB-HN2016 | | Aedes albopictus | |
| 013 | GCF_002379145.2 | Complete | wAlbB-FL2016 | | Aedes albopictus | |
| 014 | GCF_004171285.1 | Complete | wAlbB | | Aedes albopictus | |
| 015 | GCF_004795935.1 | Complete | wBm | TRS | Brugia malayi | |
| 016 | GCF_004795955.1 | Complete | wMau | | Drosophila mauritiana | |
| 017 | GCF_004795975.1 | Complete | wMau | | Drosophila mauritiana | |
| 018 | GCF_006542295.1 | Complete | wCauA | | Carposina sasakii | |
| 019 | GCF_007971685.1 | Complete | wMel_N25 | | Drosophila melanogaster | |
| 020 | GCF_007972595.1 | Complete | wMel_I23 | | Drosophila melanogaster | |
| 021 | GCF_007972745.1 | Complete | wMel_ZH26 | | Drosophila melanogaster | |
| 022 | GCF_008033215.1 | Complete | wAna | W2.1 | Drosophila ananassae | |
| 023 | GCF_008245065.1 | Complete | wMeg | | Chrysomya megacephala | |
| 024 | GCF_009732755.1 | Complete | wIrr | | Haematobia irritans | |
| 025 | GCF_012030695.1 | Complete | wBp | FR3 | Brugia pahangi | |
| 026 | GCF_012277295.1 | Complete | wCfeT | | Ctenocephalides felis | |
| 027 | GCF_012277315.1 | Complete | wCfeJ | | Ctenocephalides felis | |
| 028 | GCF_013365435.1 | Complete | wLsig | | Litomosoides sigmodontis | |
| 029 | GCF_013365455.1 | Complete | wDimm | FR3 | Dirofilaria immitis | |
| 030 | GCF_013365475.1 | Complete | wCtub | 55YT | Cruorifilaria tuberocauda | |
| 031 | GCF_013365495.1 | Complete | wDcau | 362YU | Dipetalonema caudispina | |
| 032 | GCF_013458815.1 | Complete | NA | dawsonii | Diaphorina citri | |
| 033 | GCF_016584325.1 | Complete | wMelpop | | Drosophila melanogaster | |
| 034 | GCF_016584355.1 | Complete | wMelpop2 | | Drosophila melanogaster | |
| 035 | GCF_016584375.1 | Complete | wMelOctoless | | Drosophila melanogaster | |
| 036 | GCF_016584405.1 | Complete | wMelCS_b | | Drosophila melanogaster | |
| 037 | GCF_016584425.1 | Complete | wMel | | Drosophila melanogaster | |
| 038 | GCF_017604245.1 | Complete | wCin2USA1 | | Rhagoletis cingulata | |
| 039 | GCF_017869285.1 | Complete | wWpum | MIAOwWpum | Wiebesia pumilae | |
| 040 | GCF_017896245.1 | Complete | wMel | GV_2018_4 | Aedes aegypti (transinfected) | |
| 041 | GCF_017896265.1 | Complete | wMel | GV_2018_3 | Aedes aegypti (transinfected) | |
| 042 | GCF_017896285.1 | Complete | wMel | GV_2018_6 | Aedes aegypti (transinfected) | |
| 043 | GCF_017896305.1 | Complete | wMel | GV_2018_5 | Aedes aegypti (transinfected) | |
| 044 | GCF_017896325.1 | Complete | wMel | GV_2018_7 | Aedes aegypti (transinfected) | |
| 045 | GCF_017896345.1 | Complete | wMel | GV_2018_2 | Aedes aegypti (transinfected) | |
| 046 | GCF_017896365.1 | Complete | wMel | GV_2018_1 | Aedes aegypti (transinfected) | |
| 047 | GCF_018141665.1 | Complete | NA | Spic_B | Spodoptera picta | |
| 048 | GCF_018467115.1 | Complete | wYak | KB166 | Drosophila yakuba | |
| 049 | GCF_018467135.1 | Complete | wSan | KB161 | Drosophila santomea | |
| 050 | GCF_018689955.1 | Complete | wTei | KB156 | Drosophila simulans | |
| 051 | GCF_018690035.1 | Complete | wMa | KB154 | Drosophila simulans | |

| | | | | | | |
|---|---|---|---|---|---|---|
| 052 | GCF_018690095.1 | Complete | wAu | KB177 | Drosophila simulans | |
| 053 | GCF_019665805.1 | Complete | wAlbB-Q | | Aedes aegypti (transinfected) | |
| 054 | GCA_000953315.1 | Complete | wAu | | Drosophila simulans | |
| 055 | GCA_020995475.1 | Complete | wCcep | SYLI2103 | Corcyra cephalonica | |
| 056 | GCF_000073005.1 | Chromosome | wPip | Pel (host) | Culex quinquefasciatus | |
| 057 | GCF_001439985.1 | Chromosome | wTpre | | Trichogramma pretiosum | |
| 058 | GCF_001758565.1 | Chromosome | wInc_Cu | | Drosophila incompta | |
| 059 | GCF_003999585.1 | Chromosome | wTabA | China 1 | Bemisia tabaci | |
| 060 | GCF_013096355.2 | Chromosome | wDi | KPSwDI15P40 | Diaphorina citri | |
| 061 | GCF_013096535.2 | Chromosome | wDi | KPSwDI10P38 | Diaphorina citri | |
| 062 | GCF_013096725.2 | Chromosome | wDi | KPSwDI05P26 | Diaphorina citri | |
| 063 | GCF_014107455.1 | Chromosome | wNik | | Drosophila nikananu | |
| 064 | GCF_014107475.1 | Chromosome | wStv | | Drosophila sturtevanti | |
| 065 | GCF_014771645.1 | Chromosome | wChem | PL13 | Cimex hemipterius | |
| 066 | GCF_017869155.1 | Chromosome | wCsol | MIAOwCsol | Ceratosolen solmsi | |
| 067 | GCF_000156735.1 | Scaffold | wPip | JHB (host) | Culex quinquefasciatus | |
| 068 | GCF_000174095.1 | Scaffold | wUni | | Muscidifurax uniraptor | Excluded from phylogenetic analysis because of low BUSCO score. |
| 069 | GCF_000204545.1 | Scaffold | wVitB | | Nasonia vitripennis | |
| 070 | GCF_000475015.1 | Scaffold | wMelPop | | Drosophila melanogaster | |
| 071 | GCF_000689175.1 | Scaffold | wGmm | | Glossina morsitans morsitans | |
| 072 | GCF_000723225.2 | Scaffold | wPip_Mol | | Culex molestus | |
| 073 | GCF_001266585.1 | Scaffold | NA | Ob_Wba | Operophtera brumata | |
| 074 | GCF_001675785.1 | Scaffold | wNfe | | Nomada ferruginata | |
| 075 | GCF_001752665.1 | Scaffold | wPpe | | Pratylenchus penetrans | |
| 076 | GCF_002204235.2 | Scaffold | wWb | | Wuchereria bancrofti | |
| 077 | GCF_002300525.1 | Scaffold | wSpc | | Drosophila subpulchrella | |
| 078 | GCF_004685025.1 | Scaffold | wMau | | Drosophila mauritiana | |
| 079 | GCF_005862115.1 | Scaffold | wYak_CY17C | | Drosophila yakuba | |
| 080 | GCF_005862135.1 | Scaffold | wTei_cascade_4_2 | | Drosophila teissieri | |
| 081 | GCF_007115015.1 | Scaffold | wStri | | Laodelphax striatellus | |
| 082 | GCF_007115045.1 | Scaffold | wLug | | Nilaparvata lugens | |
| 083 | GCF_009012935.1 | Scaffold | wOneA1 | | Nasonia oneida | |
| 084 | GCF_014129515.1 | Scaffold | wTri-2 | | Drosophila triauraria | |
| 085 | GCF_014129525.1 | Scaffold | wTro | | Drosophila tropicalis | |
| 086 | GCF_014129535.1 | Scaffold | wNeo | | Drosophila neotestacea | |
| 087 | GCF_014129565.1 | Scaffold | wOrie | | Drosophila orientacea | |
| 088 | GCF_014129605.1 | Scaffold | wBai | | Drosophila baimaii | |
| 089 | GCF_014129615.1 | Scaffold | wBor | | Drosophila borealis | |
| 090 | GCF_014129645.1 | Scaffold | wBic | | Drosophila bicornuta | |
| 091 | GCF_014129655.1 | Scaffold | wAra | | Drosophila arawakana | |
| 092 | GCF_014129685.1 | Scaffold | wBif | | Drosophila bifasciata | |
| 093 | GCF_014333535.1 | Scaffold | wAgra | Ano62 | Anoplolepis gracilipes | |
| 094 | GCF_014354315.1 | Scaffold | wSh | | Drosophila sechellia | |
| 095 | GCF_014354335.1 | Scaffold | wMelCS | | Drosophila melanogaster | |
| 096 | GCF_014354345.1 | Scaffold | wMel | PC75 | Drosophila melanogaster | |
| 097 | GCF_014534705.1 | Scaffold | NA | WolPenNig | Pentalonia nigronervosa | |
| 098 | GCF_017916155.1 | Scaffold | wMel | FFD25 | Drosophila melanogaster | |
| 099 | GCF_017916175.1 | Scaffold | wAur | SP11-11 | Drosophila auraria | |
| 100 | GCF_020278605.1 | Scaffold | wMpe1 | | Mansonella perstans | |
| 101 | GCF_020278625.1 | Scaffold | wMoz2 | | Mansonella ozzardi | |
| 102 | GCF_902713635.1 | Scaffold | wCobs | wCobs-BR2009-alpha | Cardiocondyla obscurior | |
| 103 | GCF_902713645.1 | Scaffold | wCobs | wCobs-JP2010-OypB | Cardiocondyla obscurior | |
| 104 | GCF_000167475.1 | Contig | wAna | | Drosophila ananassae | Excluded from phylogenetic analysis because of low BUSCO score. |
| 105 | GCF_000242415.2 | Contig | wAlbB | | Aedes albopictus | |
| 106 | GCF_000333775.1 | Contig | wBol1-b | | Hypolimnas bolina | |

| | | | | | |
|---|---|---|---|---|---|
| 107 | GCF_000333795.1 | Contig | wSuzi | valsugana | Drosophila suzukii |
| 108 | GCF_000742435.1 | Contig | wRec | | Drosophila recens |
| 109 | GCF_001027565.1 | Contig | wVulC | | Armadillidium vulgare |
| 110 | GCF_001637495.1 | Contig | wStri | | Laodelphax striatellus |
| 111 | GCF_001675695.1 | Contig | wNfla | | Nomada flava |
| 112 | GCF_001675715.1 | Contig | wNleu | | Nomada leucophthalma |
| 113 | GCF_001675775.1 | Contig | wNpa | | Nomada panzeri |
| 114 | GCF_001983615.1 | Contig | wVitA | | Nasonia vitripennis |
| 115 | GCF_001983635.1 | Contig | wUni | | Muscidifurax uniraptor |
| 116 | GCF_002318985.1 | Contig | wAus | | Plutella australiana |
| 117 | GCF_002907405.1 | Contig | wRi | | Drosophila ananassae |
| 118 | GCF_002907445.1 | Contig | wMel_AMD | | Drosophila melanogaster |
| 119 | GCF_002907525.1 | Contig | wMel_KL | | Drosophila melanogaster |
| 120 | GCF_003344345.1 | Contig | wCon | | Cylisticus convexus |
| 121 | GCF_003671365.1 | Contig | wAna_India | | Drosophila ananassae |
| 122 | GCF_003671375.1 | Contig | wAna_Indonesia | | Drosophila ananassae |
| 123 | GCF_003671405.1 | Contig | wAna_Hawaii | | Drosophila ananassae |
| 124 | GCF_004795415.1 | Contig | wAlbB | | Aedes albopictus |
| 125 | GCF_005862095.1 | Contig | wSan_Quija630.39 | | Drosophila santomea |
| 126 | GCF_006334525.1 | Contig | wLcla | GBW | Leptopilina clavipes |
| 127 | GCF_010820705.1 | Contig | wAdent | | Apterostigma dentigerum |
| 128 | GCF_013309895.1 | Contig | wApol | K3 | Atemnus politus |
| 129 | GCF_013317055.1 | Contig | wApol | K5 | Atemnus politus |
| 130 | GCF_013366805.1 | Contig | wLbra | 37PF | Litomosoides brasiliensis |
| 131 | GCF_013366855.1 | Contig | wMhie | 81YU | Madathamugadia hiepei |
| 132 | GCF_016031645.1 | Contig | wKgib | MIAO-Lgi1 | Kradibia gibbosae |
| 133 | GCF_018454445.1 | Contig | wCer5 | | Rhagoletis cerasi |
| 134 | GCF_018454455.1 | Contig | wCer4 | | Ceratitis capitata (transinfected from Rhagoletis cerasi) |
| 135 | GCF_018454475.1 | Contig | wCer1 | | Rhagoletis cerasi |
| 136 | GCF_018491625.1 | Contig | wAnM | | Anopheles moucheti |
| 137 | GCF_018491735.1 | Contig | wAnD | | Anopheles demeilloni |
| 138 | GCF_018555315.1 | Contig | wEcas | | Erebia cassioides |
| 139 | GCF_020405475.1 | Contig | wCon | | Tribolium confusum |
| 140 | GCF_900097055.1 | Contig | NA | WBT1 | Bemisia tabaci |
