## Supplementary figures and images for "Two complete genomes of male-killing *Wolbachia* infecting *Ostrinia* moth species illuminate their evolutionary dynamics and association with hosts"

### Supplemental Figures

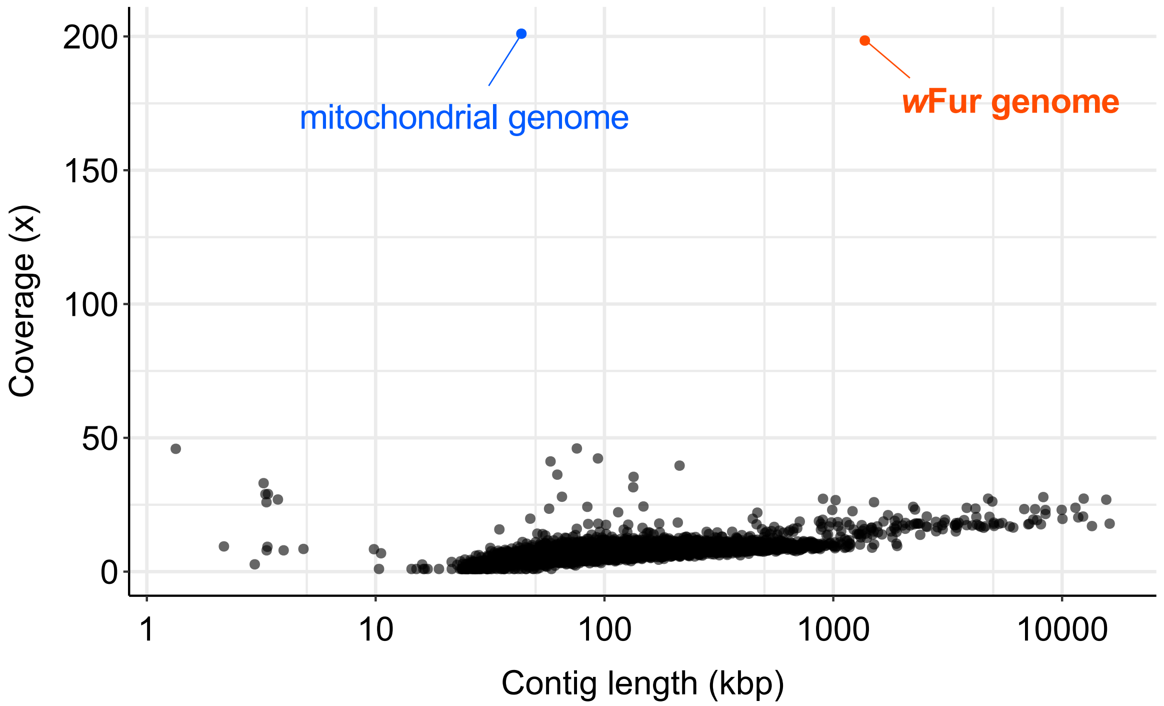

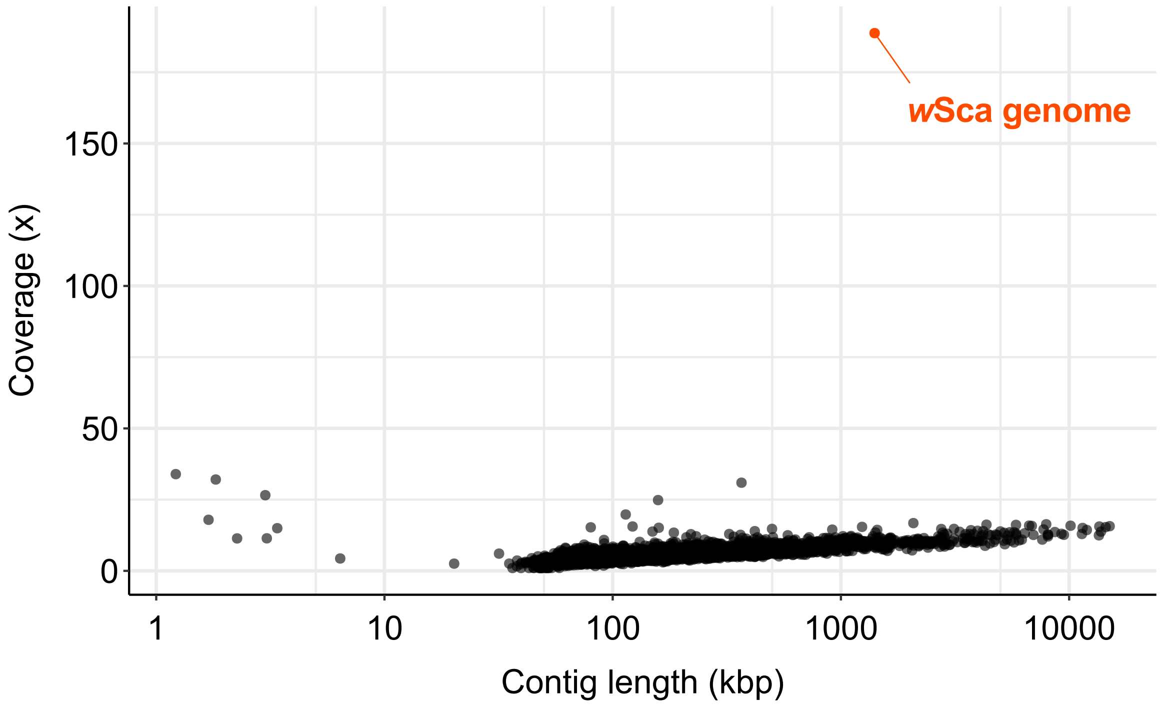

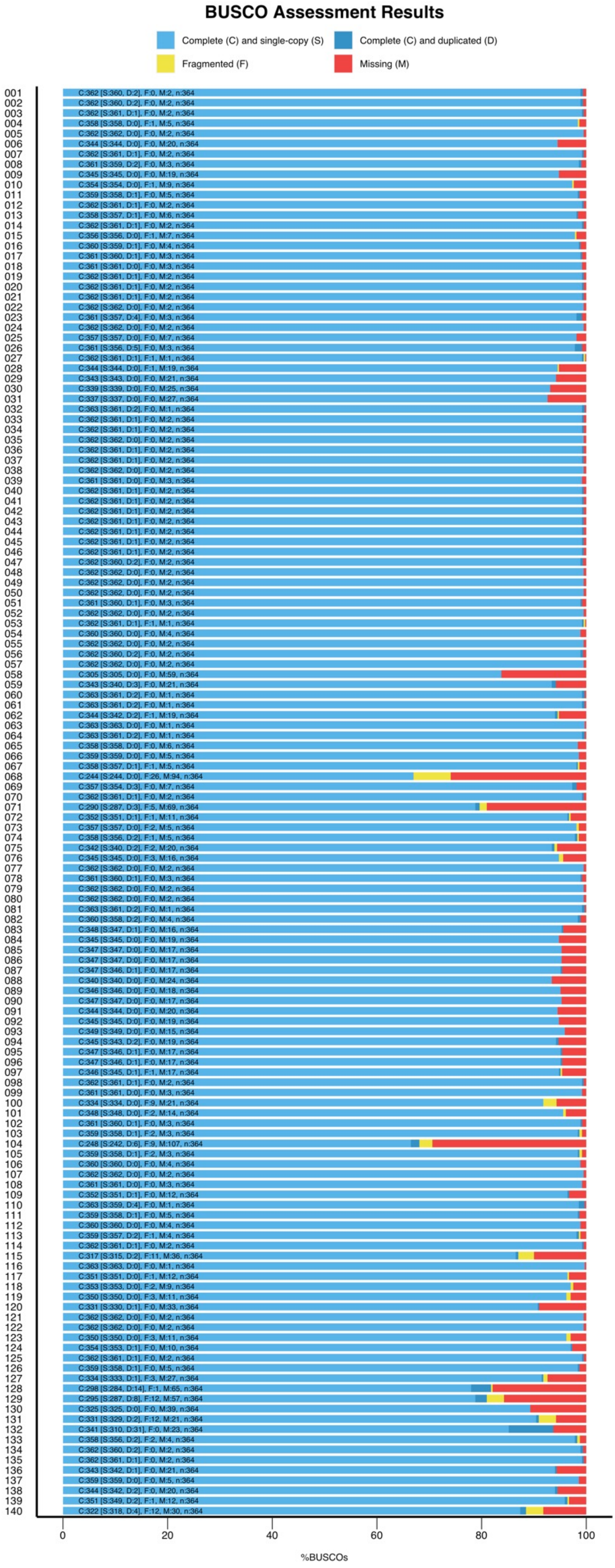

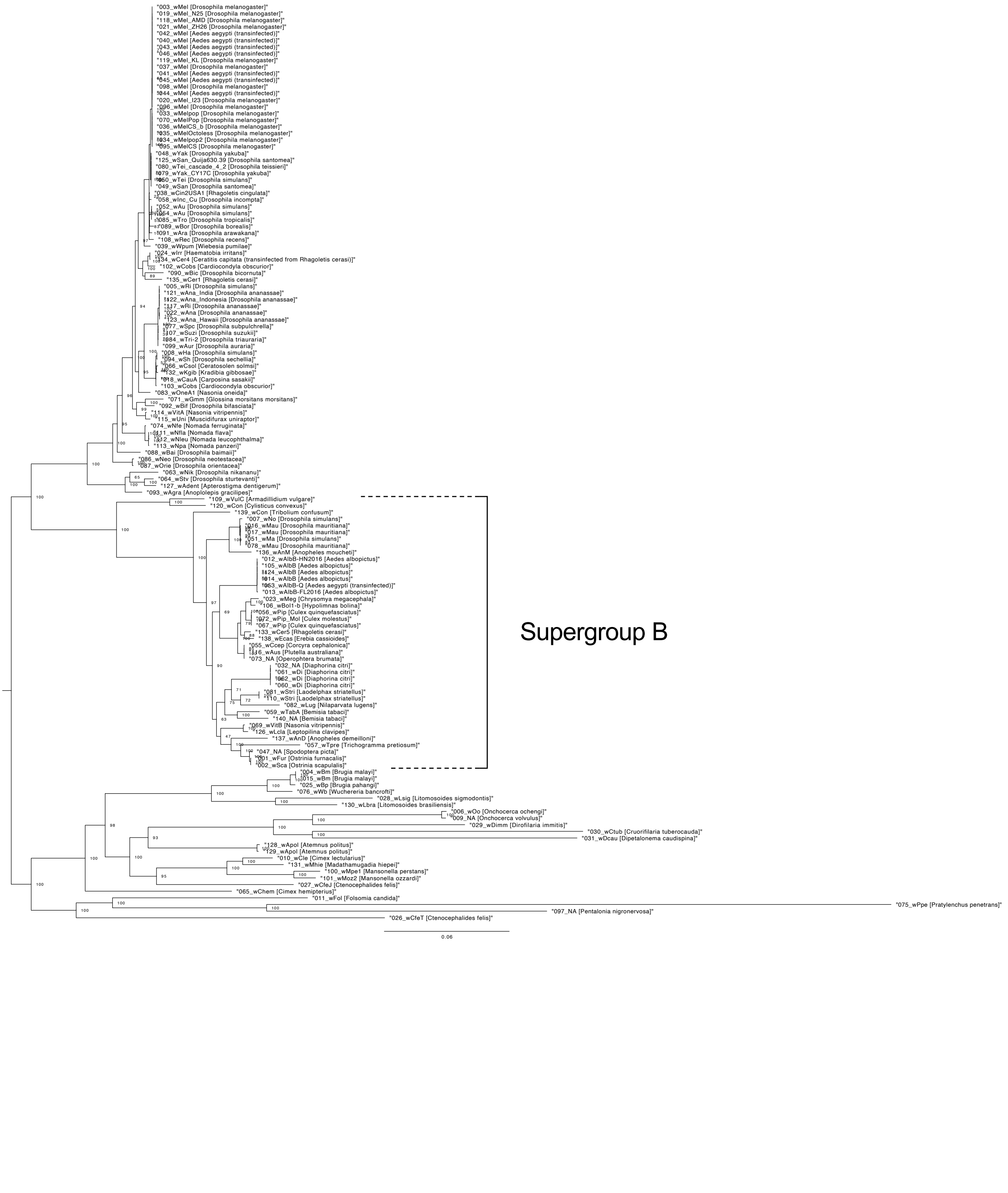

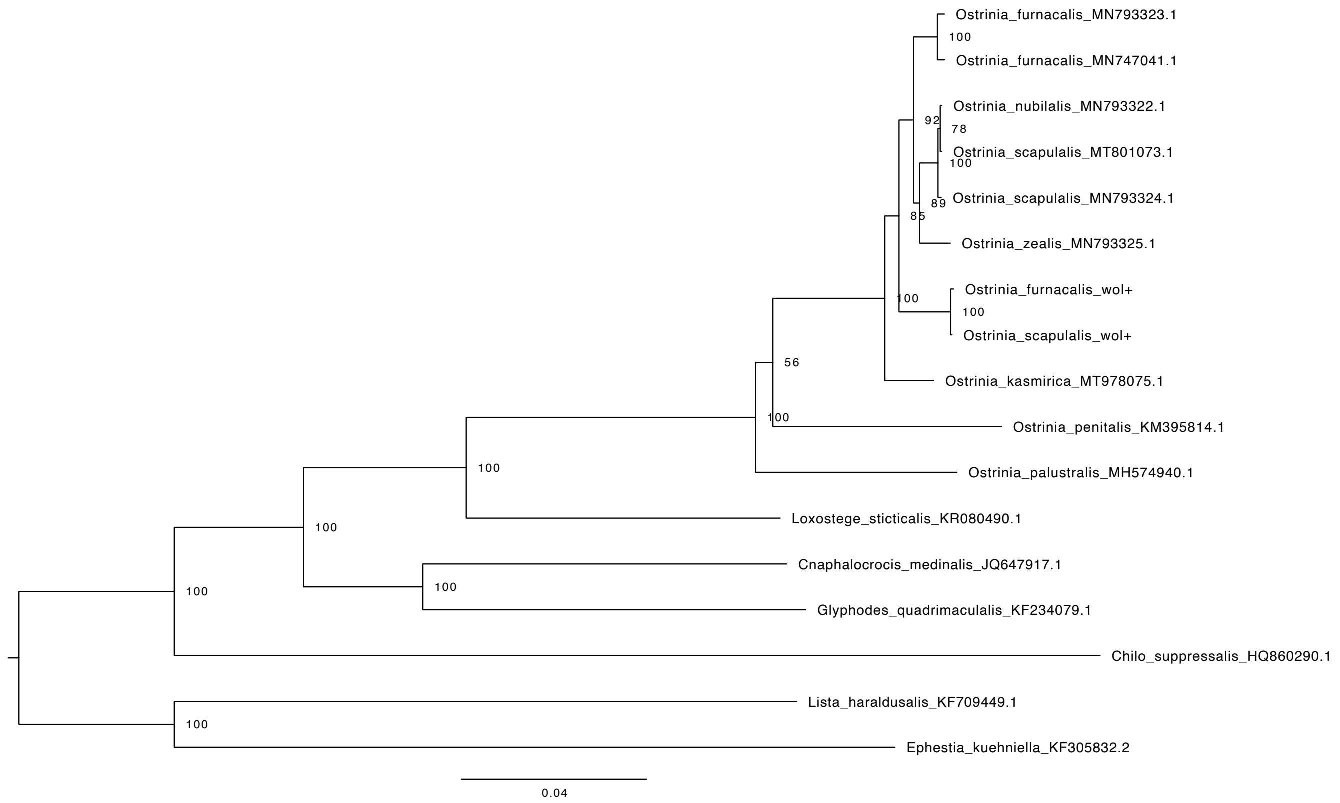
